# Perfect genetic biomarkers for cancer from a fresh view of gene dysregulation

**DOI:** 10.1101/2022.07.25.501449

**Authors:** Gabriel Gil, Augusto Gonzalez

## Abstract

Over the last decades, a host of gene expression profiles of tumor and normal tissue samples have been recorded by many microarray and RNA-Seq projects. Much of this big data awaits a full understanding and exploitation for translational cancer research. In particular, the pressing need to discover gene panels for diagnosis and therapy have not received yet a definitive answer. Here, we tackle such a question through rigorous mining of some of the currently available data. Our mining scheme rests on formal concept analysis and rough set theory and allows us to identify perfect gene panels for twelve of the solid tumors reported in the TCGA database. We dub them ‘perfect gene panels’ because they perfectly discriminate between normal and tumor samples. To wit, testing the gene expression profiles against a tumor or normal pattern provides no false positive and no false negative cases (i.e., 100% sensitivity and 100% specificity). Hence, perfect gene panels might be useful genetic markers for cancer diagnosis. Furthermore, we stress that such panels come in many flavors depending on the gene expression levels we choose as a pattern to check. For instance, there are perfect panels where a single gene over-expression signals a tumor and others where a single non-silenced gene is an indication of a tumor-free sample, just to mention two out of eight possible cases. Remarkably, some panels also suggest suitable genetic targets for therapeutic interventions, since they define normal samples by tuning the expression level of a single gene.

## Introduction

The Human Genome Project of the ‘90s opened the door to many large-scale omics catalogues (Collins, Morgan and Patrinos, 2003). By the next decade, the field experienced a further boost thanks to high-throughput microarrays, and next-generation RNA sequencing (RNA-Seq) (Chu and Corey, 2012). The new technology brought about more recent and specialized databases with a focus on biomedical applications (Haque, Engel, Teichmann and Lönnberg, 2017). A notorious example is The Cancer Genome Atlas (TCGA), which discloses potentially crucial information for cancer detection and treatment, as well as for a fundamental understanding of oncogenesis (TCGA Research Network, 2013; Hutter and Zenklusen, 2018). TCGA stores a host of gene expression profiles on tumor and normal tissue samples for 33 cancer types (TCGA Research Network, 2006). All of this data is publicly available for mining and analysis in the interest of discovering specific set of genetic markers and targets (TCGA Research Network, 2006). As expected, current TCGA analytics mirrors the experimental feat of collecting such a big data (Cheng, Dummer and Levesque, 2015; Liñares-Blanco, Pazos and Fernandez-Lozano, 2021). However, a definitive answer about the most adequate set of genes for diagnosis and therapy is yet to come.

In this context, we provide another attempt to find such a suitable set of genes. The only desideratum is that the gene panel should be able to discriminate perfectly between normal and tumor samples on the grounds of their gene expression levels. A gene panel that is up to the task is considered perfect. Our scheme is light on assumptions and, in a sense, it is closer to mining than to the analysis of gene expression profiles.

We suppose that there are only two relevant classes of expression levels: normal and tumor-like. The temptation is to ascribe a normal status to expression levels around an average value for normal samples, but this approach may rule out important genes which are known to have a fluctuating expression in a normal tissue, a behavior akin to circadian clock genes (Andreani, Itoh, Yildirim, Hwangbo, and Allada, 2015), for example. Therefore, starting from expression levels around, above and below a given value, we define all the possible setups in which the putative normal and tumor-like expression classes may come about.

Our core procedure is based on formal concept analysis (FCA) and rough set theory (RST), both with a growing number of applications in omics (for FCA: Gebert, Motameny, Faigle, Forst, Schrader, 2008; Choi, Huang, Lam, Potter, Laubenbacher, and Duca, 2008; Motameny, Versmold, Schmutzler, 2008; Amin, Kassim, Hassanien and Hefny, 2012; Kaytoue-Uberall, Duplessis, Napoli, 2008; Kaytoue, Kuznetsov, Napoli, Duplessis, 2011; for RST: Maji and Paul, 2011; Midelfart, Komorowski, Nørsett, Yadetie, Sandvik, Lægreid, 2002; Dai and Xu, 2013; Li and Zhang, 2006; Mishra, Dash, Rath, and Acharya, 2011; Pati, Das and Ghosh, 2013). The main scope of these techniques is to discover patterns (namely, formal concepts or rough sets) in multivariate data, where a set of attributes is made to correspond to a set of objects by means of a specific relation (Duntsch and Gediga, 2002; Lai and Zhang, 2009). Notice that this is exactly the framework under consideration, provided the following mapping: genes in place of attributes, clinical samples instead of objects and gene expression profiles manifesting the relation between them (e.g., Motameny, Versmold, Schmutzler, 2008).

Finding formal concepts and rough sets is a non-trivial task, *a fortiori* when the data at hand is so big as in the case of TCGA (hundreds of gene expression profiles per cancer type, 60483 gene expression values registered). Here, the formal concepts of interest are characterized by a set of genes with either tumor-like expression levels in all tumor samples (tumor concepts), or normal-like expression levels in all normal samples (normal concepts). Selecting genes on such grounds may still lead to imperfect discrimination given that we are not constraining the expression levels for the complementary group of samples. However, for various setups of putatively normal and tumor-like expression levels we disclose perfect gene panels in this fashion. To wit, at least one deviation from normal-like expression levels for normal concept genes correspond to a tumor sample, and vice versa. Therefore, our perfect gene panels define both a formal concept for a group of samples (either normal or tumorous) and a rough-set concept for the complementary group (of tumor or normal samples, respectively). We stress that perfect gene panels offer clear-cut classifications with no false positives and no false negatives (i.e., 100% sensitivity and 100% specificity).

Albeit perfect, in many cases the pool of genes (stemming from tumor or normal concepts alike), is extremely large (with up to ~32000 genes), ruling them out for any practical application. Remarkably, following a simple heuristic, we are able to shrink out these large pools to small panels (known as reducts in RST), with no more than 20 genes. These perfect gene panels might be amenable biomarkers for cancer diagnosis.

When the perfect gene panel is built up from the tumor concept, any normal sample can be defined as such by a single gene deviating from the tumor-like expression level. Such a behavior suggests that by altering a single gene expression of a tumor concept in a tumor cell, one may render it non-tumorous, either by reverting it to its previously normal status or by causing it to perish. Anyhow, this sort of genes might be suitable targets for therapy.

Perfect gene panels come in many flavors. First, there are those related to tumor and normal concepts. In turn, there is a variety of tumor and normal concepts stemming from different setups for the tumor and normal-like expression levels. Some of these panels have a clear interpretation in terms of the state-of-the-art taxonomy of driver genes. For instance, there are perfect panels where a single gene over-expression signals a tumor. Those might be labeled oncogenes, provided an interventional proof of their causal power over cancer. Conversely, for other perfect panels, a single non-silenced gene is an indication of a tumor-free sample. With the previous causal caveat, such a description fits our current understanding of tumor suppressors. Other panels have a less straightforward interpretation, possibly involving cooperative tumor suppressors, oncogenes and genes with oscillating expression levels.

We explore twelve solid tumors among the 33 cancer types in TCGA. Having more than 20 normal samples, the selected cases were reasonable candidates to discover gene expression patterns which justify clear-cut classifications. For all tissues under study, we find perfect gene panels with possible applications in diagnosis and therapy.

In the Results, we illustrate our workflow for a case in point (namely, prostate adenocarcinoma). Our findings for the rest of the selected cancer types are also summarized therein. The following Discussion is mainly devoted to the gene dysregulation as it is understood in this paper and how it allows us to better describe homeostasis and cancer, however it also covers possible application of such panels and their role in tumorigenesis. To close the section, we offer an outlook for future translational research in the same lines. The Supplementary Information (SI) includes, first and foremost, an inventory of gene panels that perfectly discriminate between normal and tumor samples for every cancer type under study. A detailed account of our methodology and a link to the online repository hosting our code can also be found within the SI. Additionally, we provide an analysis of the robustness of our gene panels, particularly in light of a different universe of clinical samples.

## Results

We select 12 cancer types for a systematic study (see Table I). All of the chosen cancers manifest as solid tumors, particularly in the liver, breast, colon, head and neck, kidneys, lungs, prostate, stomach, thyroids and uterus. We are including five (out of six) of the most common cancer types (breast, lung, colon, prostate and stomach), with an incidence over a million cases in 2020, together with the most common causes of cancer deaths (lung, colon, liver, stomach and breast), with over half a million in 2020 (worldwide statistics reported by WHO (2021)).

**Table I.** Cancer types under study, along with their TCGA abbreviation.

| Cancer types | Abbreviation |
| --- | --- |
| Liver hepatocellular carcinoma | LIHC |
| Breast invasive carcinoma | BRCA |
| Colon adenocarcinoma | COAD |
| Head and Neck squamous cell carcinoma | HNSC |
| Kidney renal clear cell carcinoma | KIRC |
| Kidney renal papillary cell carcinoma | KIRP |
| Lung adenocarcinoma | LUAD |
| Lung squamous cell carcinoma | LUSC |
| Prostate adenocarcinoma | PRAD |
| Stomach adenocarcinoma | STAD |
| Thyroid carcinoma | THCA |
| Uterine Corpus Endometrial Carcinoma | UCEC |

TCGA RNA-Seq profiles are associated with biopsies performed on cancer patients (TCGA Research Network, 2006). Some of the clinical samples are normal (healthy) and other tumorous, and they are classified accordingly by histopathological procedures (TCGA Research Network, 2013). Being the focus on cancer patients, data is naturally biased towards tumor samples and, in fact, there are some tissues for which no normal sample is available in database. Importantly, we need both classes of samples to be well represented within the data universe [SI:C1] to accomplish our goal, which is to provide a sound discrimination between normal and tumor tissue on the grounds of their gene expression profiles. The selection of cancer types for our systematic study is precisely motivated by the number of normal samples within the data. For the cases under study, TCGA reports on more than 20 normal samples per cancer type.

For each tissue, we take the log fold change of the gene expression with respect to its normal reference (i.e., an average over normal samples). This sort of quantity places up and down regulation on equal footing in the following analysis. For constitutive genes, we expect the log fold change to distribute normally (in a Gaussian-like fashion) around a reference for normal (non-tumorous) samples. Yet, this is but a possibility among a variety of distributions that come about depending on the gene and its role in homeostasis. On the other hand, we know that a tumor shifts sensibly the expression levels for many genes. In particular, we expect cancer to be driven by an increase or decrease in the expression of oncogenes or tumor suppressors, respectively. However, there is a wide range of tumor dysregulation patterns taking place for other kinds of genes.

We focus on PRAD to illustrate the previous remarks. In Figure 1, we plot smooth kernel distributions of the gene expression in tumor and normal tissue for a selection of genes [SI:C2]. We stress the diversity of non-trivial shapes of both tumor and normal distributions and the qualitative differences among them. Figure 1 shows normal expression values distributed in an asymmetric and multimodal way. For instance, normal distributions for *SEPTIN10, EPHA10, LYPD3, PRRT3-AS1* and *SNHG3* are bimodal, suggesting an oscillatory and in some cases (e.g., *SNHG3*) pulsatile gene expression [SI:C3]. The tumor dysregulates the latter genes by arresting their oscillation (especially in *SEPTIN10* and *LYPD3*), and in the case of *EPHA10, PRRT3-AS1* and *SNHG3*, by tuning up their mean expression level. Such a halted oscillation is consistent with an alteration in the expression of transcription regulators (for example, a defect in the activators for *SEPTIN10* and in repressors for *LYPD3*). Granted that the normal-to-tumor transition may not have a clear interpretation in other cases of Figure 1, a strong reorganization of expression is also apparent for the rest of the genes.

**Figure 1:**
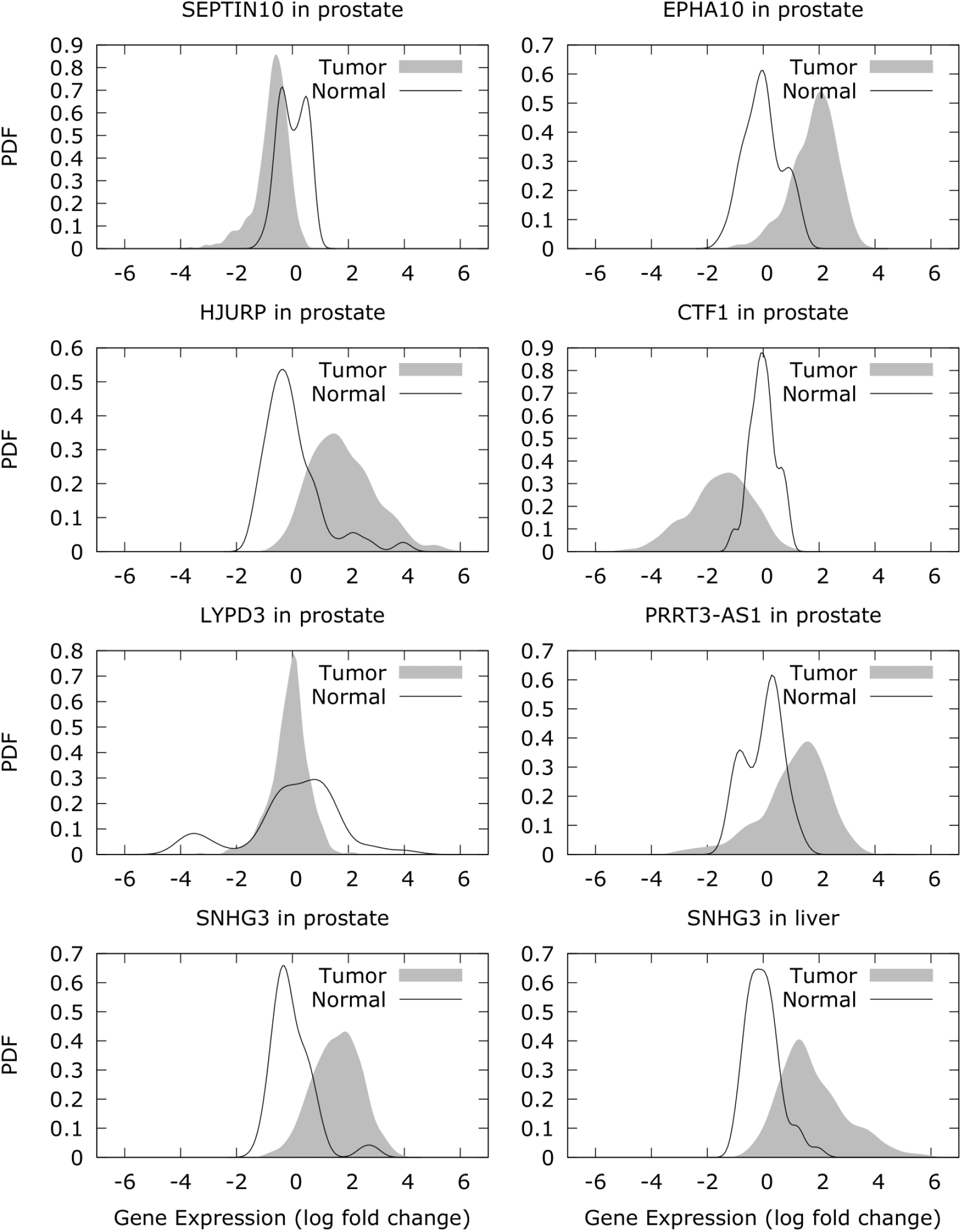
Probability density functions (PDFs) of normal and tumor gene expression in prostate and liver for a set of representative classifier genes. In the abscissa, we have the log fold change of the expression with respect to its geometric mean in normal samples. Data is obtained from TCGA, and Kernel Density Estimation is performed using Gaussians and the Silverman rule for the choice of the bandwidth.

In fact, there is a common feature underlying all cases in Figure 1. For each of them, there are expression windows where normal and tumor distributions do not overlap. In detail, such genes exhibit expression ranges deprived of either tumor or normal samples, while covering part of the complementary group. For instance, *EPHA10* is only expressed above 2.5 times the reference level for tumors, and *HJURP* is only expressed below half the reference value for normal samples. As expected, the aforementioned expression windows depend on the tissue. In the prostate, *SNHG3* expression about 2-6 times the normal reference value indicates a tumor, whereas in the liver, it needs to be overexpressed beyond ~4 times the reference level to arrive at the same conclusion (see last two panels of Figure 1).

A classification based on any such genes yields no type I errors, provided the null hypothesis to be that the sample is of the only kind excluded from a specific expression window [SI:C4]. Assessing whether a gene expression lie within such exclusive zone [SI:C5] or not amounts to reject or accept the null hypothesis of it being a tumor or a normal sample, depending on the case. To select significant genes in this category genome-wide, we constrained the probability of generating the classification of samples by chance with a p-value < 1% [SI:C6]. In the average cancer type, we find 39% of significant genes (out of ~60000).

Classifier genes can be grouped into two subclasses, depending on whether the classification they suggest is *a priori* free from false normal samples (e.g., *HJURP*) or free from false tumors (e.g., *CTF1*). Some genes belong to both subgroups since they feature an expression windows without tumors, and another without normal samples. For example, *EPHA10* expression in the prostate is exclusive of normal samples below nearly half of the reference level, and exclusive of tumors above ~2.5 times it. For classifier genes without false tumors, expression levels beyond normal identify some of the tumor instances (namely, tumor detection is 100% specific), whereas for classifier genes lacking false normal samples, it is the departure from tumor expression levels which allows us to pinpoint some of the normal cases (namely, normal detection is 100% specific).

In our approach, a normal-like gene expression is either specific of normal samples or not specific of tumors, whereas a tumor-like expression is specific of tumors or not specific of normal samples (see Discussion). We define a normal concept as the largest set of genes taking normal-like expression values for all normal samples (i.e., the class of classifier genes without false tumors). By the same token, a tumor concept is the largest set of genes taking tumor-like expression values for all tumors (i.e., the class of classifier genes without false normal samples).

In the average cancer type, nearly 2.9% of the genes belong to the tumor concept, suggesting that concurrent dysregulation is not so widely spread across the genome. The fact that more than one third of the genome and the vast majority of classifier genes fall within the normal concept is, on the one hand, consistent with the picture of cancer as a highly entropic state of the gene regulatory network, and on the other, an indication of the large amount of potential genetic triggers of cancer (see Discussion).

Taken independently, genes described above still misclassify some of the samples due to type II errors [SI:C7]. For instance, *CFT1* from Fig. 1 cannot discriminate between tumor and normal tissue above one fourth of its reference expression level. However, using combinations of classifier genes of the same kind we may reduce type II errors. Indeed, accepting the null hypothesis for every classifier gene in a collection is a more stringent criterion of acceptance for the null hypothesis. Therefore, following such a scheme we expect to maximize the number of rejections without it being detrimental to the fact that they should be *true* rejections. This procedure leads not only to the reduction but to the complete elimination of type II errors (and hence, to a perfect classification of samples) for the twelve tissues here considered and either kind of classifier genes [SI:C8]. For instance, by assembling *CFT1* with other 14 genes of the same kind in the same panel (see SI) we classify all prostate samples into normal and tumor without mistakes.

We use ‘perfect panel’ to denote the reduced set of genes that guarantee a faultless classification. Perfect panels drawn from the normal concept define a minimal expression pattern which is exclusive of the homeostatic state [SI:C9]. Hence, this is the panel of cancer drivers, including both oncogenes and tumor suppressors. Indeed, a tumor-like expression of any such genes indicates a tumor and therefore (with the usual covariation-is-not-causation caveat), can be said to trigger tumorigenesis (see Discussion). Among the 7 members of the normal-concept panel for the prostate, we get *COL10A1*, pertaining to the pathway of Molecular Mechanisms of Cancer, and *EPHA10*, having p53 as a transcription factor and coming from a family of genes that is implicated in cancer development and progression (Stelzer et al., 2016).

On the other hand, perfect panels drawn from the tumor concept define a minimal expression pattern which is exclusive of the cancer state. Therefore, we expect cancer hallmark genes to be comprised within this panel [SI:C10]. Among the 11 genes in the prostate tumor-concept panel, we get *HJURP*, a gene related to DNA Damage and Chromosome Maintenance pathways which has been previously linked to breast and lung cancer, *SEPTIN10*, encoding a filament-forming cytoskeletal GTPase which is abundantly expressed in various tumor cell lines, *TBC1D7*, which participate in the regulation of cellular growth and differentiation, *TES*, playing a role in cell adhesion, spreading and proliferation, as well as in the inhibition tumor cell growth, and *SEPTIN5*, a translocation of which (involving also the *MLL* gene) has been reported in patients with acute myeloid leukemia (Stelzer et al., 2016). Furthermore, two of the panel genes (namely, *SEPTIN5* and *SEPTIN10)* are responsible for the top-ranked (p-value = 6.9×10^-4) enrichment of ‘cytoskeleton-dependent cytokinesis’ term in the gene ontology of biological processes (see Figure 2) (Ma’ayan lab, 2021; Xie et al., 2021). Notice that the previous enrichment analysis was performed with a small gene set (only eleven), making a two-hit result a very significant one. A comprehensive gene ontology and enrichment analysis of perfect panels corresponding to other tissues is beyond our scope. However, we stress that both normal and tumor concepts were found for all tissues under study.

**Figure 2:**
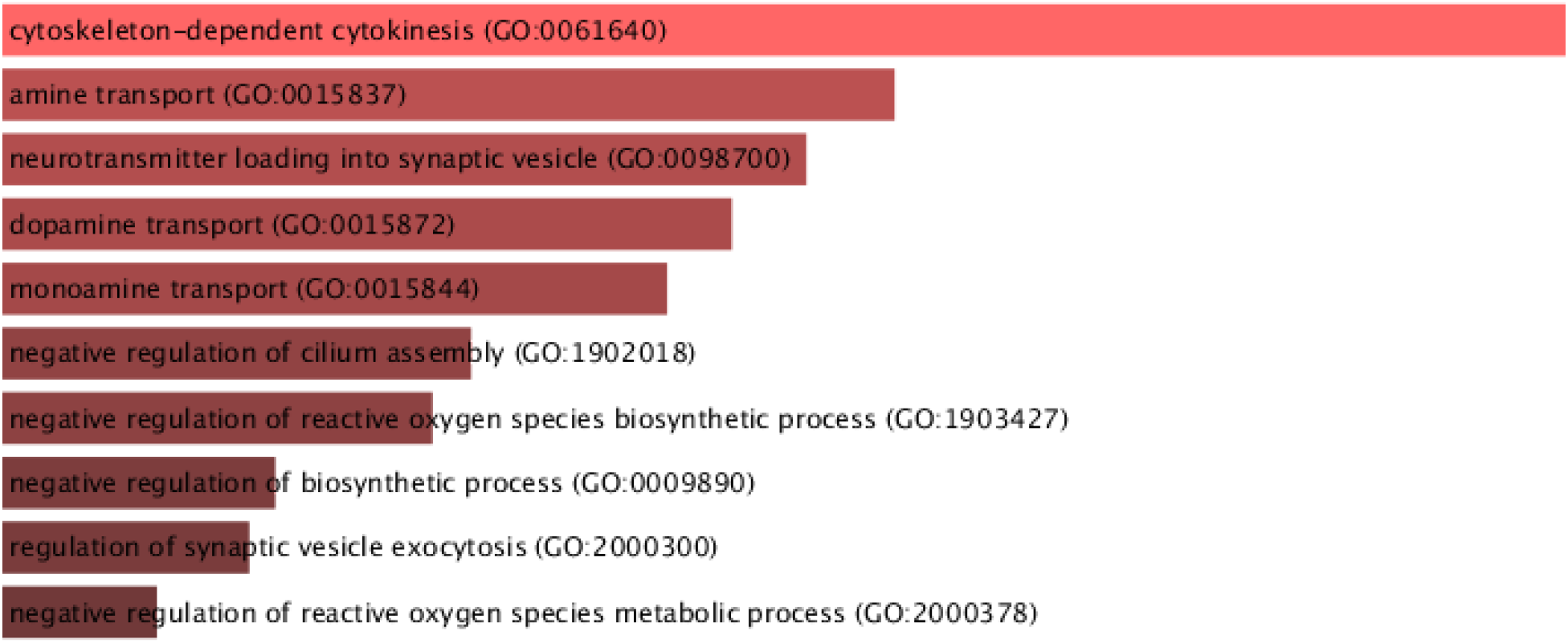
Bar chart summarizing the results of an enrichment analysis of the biological processes connected to the prostate tumor-concept panel. Associated p-values ranges from 6.9×10^-4 at the top to 1.3×10^-2 at the bottom. The 11 genes involved were *HJURP, SEPTIN10, AL512363.1, CA14, FGD5P1, SEPTIN5, TES, SLC18A2, SNORD104, TBC1D7* and *RNA5SP342*. The figure was produced by Enrichr (Ma’ayan lab, 2021).

Hitherto, we found perfect panels irrespective of whether their gene members exhibit specific sorts of dysregulation upon the transition from a normal tissue to cancer. In fact, the prostate tumor-concept panel include genes which are dysregulated in different ways. In particular, we find that both the down regulation of *HJURP* expression (below half its reference level) and the up-regulation of *TES* (above the double of its reference level) are signatures of a homeostatic control in the normal prostate. Note that this are prototypical threshold levels for considering disease-related differential expression. However, we observe that overcoming such thresholds is indicative of a healthy tissue (see Discussion).

Genes sharing the character of their dysregulation in the crossover to cancer can also be assembled into perfect panels. To this aim, we further filter classifier genes into those that are only expressed above a threshold value, below it, outside an interval, or within it for a single group of samples -be it normal or tumorous. This operation yields four normal and four tumor concepts per tissue, besides the normal and tumor concepts previously obtained from the unfiltered mixture of classifier genes.

To improve readability, we introduce shorthand names for gene sets that are suggestive of the classificatory potential of their individual members. Consider *x* to be a type of sample, whether normal or tumor. Genes that are *only* expressed *above* or *below* a threshold level for type *x* samples are referred to as ‘only *x* above’ or ‘only *x* below’, respectively. Likewise, genes that are *only* expressed *outside* or *inside* an interval for type *x* samples are called ‘only *x* outside’ or ‘only *x* inside’. ‘Only *x* in zone’ applies to genes featuring expression windows *only* visited by type *x* samples (i.e., to classifier genes without false *x* outcomes). As anticipated, the latter gene set includes the former four.

Note that panels with ‘only tumor above’ and ‘only tumor below’ must include oncogenes and tumor suppressors, in that order. As shown in Table II, every tissue features perfect panels of both of the previous kinds, as well as the ones with ‘only tumor outside’ and ‘only tumor inside’. The fact that all these sorts of perfect normal-concept panels appear together for any of the tissues under study suggests that different sorts of cancer triggers are causally connected to each other.

**Table II.** Summary of classifier genes per tissue. Each column identifies a cancer type in the TCGA terminology (see Table I). Each row represents a different set of classifier genes (see main text for shorthand notation). To distinguish normal from tumor concepts, we put the formers in shaded rows. Within each cell, we show the minimal number of genes that classify the largest amount of the samples together with the total number of genes of the same sort. A star marks the minimal gene sets that constitute perfect panels.

|  | *LIHC* | *BRCA* | *COAD* | *HNSC* | *KIRC* | *KIRP* | *LUAD* | *LUSC* | *PRAD* | *STAD* | *THCA* | *UCEC* |
| --- | --- | --- | --- | --- | --- | --- | --- | --- | --- | --- | --- | --- |
| *Only tumor above* | 3/23986* | 6/15361* | 2/17536* | 4/13293* | 4/22654* | 3/11447* | 3/20274* | 2/19596* | 8/8093* | 3/13773* | 5/5744* | 1/7825* |
| *Only normal above* | 11/40 | 10/739* | 1/876* | 8/1903* | 3/780* | 1/1140* | 5/613* | 3/1198* | 14/1415* | 5/1244* | 11/794* | 1/993* |
| *Only tumor below* | 5/3812* | 6/6701* | 1/8418* | 5/2093* | 3/9132* | 1/10263* | 4/8285* | 2/9404* | 15/3865* | 5/1499* | 6/5376* | 1/7443* |
| *Only normal below* | 5/1246* | 12/682 | 2/297* | 6/1339 | 8/191 | 5/214 | 8/449 | 3/985* | 15/915 | 5/2536* | 17/92 | 1/506* |
| *Only tumor outside* | 4/7693* | 8/8433* | 3/11107* | 6/5047* | 4/11220* | 3/13058* | 3/11487* | 3/9668* | 10/6532* | 6/3353* | 5/6216* | 3/7556* |
| *Only normal outside* | 1/1 | 1/1 | -/0 | 9/20 | -/0 | -/0 | -/0 | -/0 | 3/3 | 6/41 | 3/3 | 1/1 |

**Table II.** (cont.) Summary of classifier genes per tissue.
|  | *LIHC* | *BRCA* | *COAD* | *HNSC* | *KIRC* | *KIRP* | *LUAD* | *LUSC* | *PRAD* | *STAD* | *THCA* | *UCEC* |
| --- | --- | --- | --- | --- | --- | --- | --- | --- | --- | --- | --- | --- |
| *Only tumor inside* | 6/2803* | 19/336* | 3/1991* | 5/4049* | 7/942* | 6/2475* | 6/2355* | 8/2595* | 8/2905* | 5/6026* | 5/2062* | 4/2903* |
| *Only normal inside* | 1/1 | -/0 | -/0 | -/0 | 5/5 | 12/21 | 2/2 | 4/5 | -/0 | 1/1 | 1/1 | -/0 |
| *Only tumor in zone* | 3/27814* | 4/21949* | 1/27069* | 4/19398* | 3/32218* | 1/23893* | 2/28031* | 2/29731* | 7/15837* | 3/21122* | 4/13535* | 1/19044* |
| *Only normal in zone* | 5/1287* | 10/1420* | 1/1173* | 6/3222* | 3/975* | 1/1375* | 5/1064* | 2/2188* | 11/2327* | 4/3739* | 9/888* | 1/1498* |

On the other hand, none of the analyzed filters allows us to find perfect tumor-concept panels for all of the tissues (see Table II). Additionally, no tissue has perfect panels with ‘only normal outside’ or ‘only normal inside’. Perfect panels with ‘only normal above’ or ‘only normal below’ figure in one tissue or another, however unsystematically (see Table II). To wit, BRCA, HNSC, KIRC, KIRP, LUAD, PRAD and THCA contain only the former, UCEC, COAD, LUSC and STAD contain both, and LIHC contains only the latter. Furthermore, there are some cancer types that can be completely characterized by a single gene (see SI). That is the case of COAD with *SCARA5*, KIRP with *UMOD*, and UCEC with either *PLSCR4* or *TBC1D7*. Remarkably, microarray readings from Khamas et al. (2012) demonstrate the perfect classification character of *SCARA5* (data accessible at NCBI GEO database (Barret et al., 2013), accession GDS4382), and in fact, it has been pinpointed as a biomarker of colorectal cancer (Liu, Zheng, Shi, Cao, Zhang and Xie, 2020).

Despite there being no perfect panels of a given kind, genes of the that same kind may feature in the comprehensive panels with ‘only *x* in zone’. Take the case of LIHC, for example. It lacks a perfect panel with ‘only normal above’; however, a gene from that class (namely, *MARCO*) is included in the panel with ‘only normal in zone’.

## Discussion

### Gene dysregulation

Gene dysregulation can promote cancer (Bradner, Hnisz and Young, 2017). In fact, the discovery of genes that are relevant to carcinogenesis and tumor progression is partially guided by the assessment of gene dysregulation instances against some criteria for statistical and biological importance (Li, Dai, Liu, Sang, Li and Li, 2020). The paradigmatic case of gene dysregulation is differential expression (Ali, Lung, Sholl, Gad, Bustamante, Ali, Rhim, Deep, Zhang and Elmageed, 2018). To our purpose, that is the case where a gene gets differently expressed in tumors with respect to a normal tissue.

In the traditional view, such differential expression is relevant for cancer only if it involves a strong departure from the normal expression, it comes up systematically as either above or below normal expression levels, and it is a feature of most tumors (Li, Dai, Liu, Sang, Li and Li, 2021). A common practice is to set lower and upper bounds for normal gene expression to half and twice a reference level, respectively (Dalman, Deeter, Nimishakavi and Duan, 2012). Therefore, a gene *can be* differentially expressed only if the majority of tumors expressed it more than twice or less than half the reference. In turn, any such gene *is* differentially expressed when its expression level goes beyond the single bound over or under which most tumors express it.

Thus understood, differential expression can be a manifestation of a dysregulation, but it need not to be so. Furthermore, there are clear instances of differential expression that are not covered by the previous definition. Finally, dysregulation is by no means limited to differential expression, however broadly we may understand the term.

To illustrate the last points, let us turn back to PRAD (see SI). For such a case, the mean expression threshold that corresponds to the panel with ‘only tumor above’ is close to doubling the reference level, whereas for the panel with ‘only tumor below’, the mean expression threshold almost halves the reference. These results seem in line with the view of cancer-correlated dysregulation as differential expression. After all, genes that are comprised in our panels with ‘only tumor above/below/outside’ turns out to have the capacity to signal tumors whenever they are differentially expressed beyond prototypical bounds.

However, in our approach, the issue of differential expression cannot be settled without specifying a gene. What counts as a differential expression for a gene may not count as differential expression for another. Take the case of *EPHA10* and *UBXN6*, both from the panel with ‘only tumor above’, as an example. *EPHA10* is differentially expressed over 2.53 times its reference level in 73% of prostate tumor samples, whereas *UBXN6* is differentially expressed over 1.34 times its reference level in 18% of prostate tumor samples. Moreover, a mixture of normal and tumor samples expressed *EPHA10* and *UBXN6* below their respective and aforementioned thresholds. Therefore, an expression of *EPHA10* between twice and 2.53 times its reference cannot identify univocally a tumor, even though it would be typically mistaken as a case of a differential expression. On the other hand, an expression of *UBXN6* between 1.34 times and twice its reference can only point to a tumor, despite the fact that it is framed as normal in the standard account.

Normal gene expression is finely tuned as a result of highly specific interactions between the gene, numerous gene regulators and environmental cues. Such a detailed control is crucial for homeostasis, and therefore, ought to be reflected in a proper definition of differential expression. In particular, the definition must keep differential expression as *a feature of and only of* tumors. Conversely, notice that we do not need to demand that the differential expression for *any* gene be a common feature of *all* or *most* of the tumors, since a tumor instance may be characterized by a gene expression that is peculiar just of it or a specific subset of tumors it belongs to (Mezlini, Das and Goldenberg, 2021).

Indeed, we take differential expression to be gene-dependent and not fixed by arbitrary choices. The sole requisite we impose to determine differential expression thresholds is that of leaving an exclusion zone for normal samples in a particular gene expression axis. Evidently, such definition is not independent from the data universe. Therefore, we must only keep track of the differential expressions exhibited by genes that allow statistically significant classifications of tumors. To that aim, we ask that the exclusion zone is visited frequently enough by tumor samples, so that we have 99% confidence that such a void of normal samples does not arise by chance (just to keep a conventional p-value of 1%). These conditions leave room for violations of the customary differential expression. That is why for us *UBXN6* is differentially expressed in a number of tumor samples which is far from the majority for an expression threshold which is well below twofold the reference level (see above).

In addition, there are other two classes of differential expression that are not usually contemplated. Take the case of *PRRT3-AS1* from the panel with ‘only tumor outside’ and *SNHG3* from the panel with ‘only tumor inside’ as further examples. *PRRT3-AS1* is differentially expressed outside the range from 0.41 to 2.6 times the reference expression level in 53% of prostate tumor samples, whereas *SNHG3* is differentially expressed inside the range from 2.09 to 5.77 times the reference expression level in 60% of prostate tumor samples. Again, the given intervals are maximally chosen so as that they tightly exclude normal samples. Over and under-expressions of *PRRT3-AS1* are both symptomatic of cancer. Instead, for *SNHG3*, it is an expression within specific bounds what signals a tumor. As far as we can tell, *PRRT3-AS1* and *SNHG3* could be triggers of prostate cancer since a differential expression in either case entails the presence of a prostate tumor. Still, neither of them accommodates to the current understanding of driver genes in terms of tumor suppressors and oncogenes. This raises the question of whether it is necessary that cancer is driven by tumor suppressors or oncogenes exclusively or not.

We have claimed from the outset that dysregulation is a more general phenomenon than differential expression. There are some sorts of dysregulation that do not conform either with our definition of differential expression or the one that is more commonly held in the field. Plainly, non-differential dysregulations (as we may call them) are distinctive in that they are not manifested as alterations of gene expression levels. Therefore, they are often overlooked in the analysis of gene expression data.

In our scheme, non-differential dysregulation comes about in genes that are members of the panels with ‘only normal above’, ‘only normal below’, ‘only normal outside’, ‘only normal inside’ or ‘Only normal in zone’. Take *LYPD3* from the panel with ‘only normal outside’ as a case in point. Note that all tumors express *LYPD3* within 0.1 and 5.59 times the reference level, whereas normal expression for that same gene ranges from 0.07 to 16 times the reference level without significant exclusion windows. So far as *LYPD3* is concerned, tumor instances are literally within normal bounds. Nonetheless, *LYPD3* provide us with a secure signal of a non-tumorous sample whenever its expression level departs from the narrower tumor slot. In particular, 15% of the normal samples can be found on these grounds.

Not every non-differential dysregulation has a classificatory potential. We are only interested in those that are classificatory. To account for them, we take the same approach adopted for differential expression and swap the role of tumor and normal samples. That is, we find a maximally wide exclusion zone for tumors in a particular gene expression axis, and again, we demand that normal samples visit the exclusion zone often enough so that we are confident that such void of tumors being inexplicable by chance.

A non-differential dysregulation is in principle indistinguishable from a case of normal expression. Such is the reason why our classification scheme for non-differentially dysregulated genes, whenever attainable, grounds on detecting normal instead of tumor samples. But then, what is the nature of non-differential dysregulations? To propose a possible answer, let us explore further the case of *LYPD3* in prostate. Recall that *LYPD3* expression values in between 0.1 and 5.59 times the reference level are shared by tumor and normal samples (see Figure 1). Beyond such constraints, expression values are inaccessible to tumors but not to normal samples. However, the average and modal expression for normal and tumor samples roughly coincide. This suggests that, in the transition from normal to tumor, the gene regulatory network loses its ability to up and down regulate *LYPD3* as a result of specific affectations in the functioning of its transcription regulators.

We illustrate this point by appealing to the *Lac* operon in *E. Coli* (Alberts, Johnson, Raff, Roberts and Walter, 2002). *Lac* transcription is controlled by an activator and a repressor, acting in opposite ways. The activator enables transcription, in the absence of glucose, whereas the repressor prevents it, in the absence of lactose. As a consequence, *Lac* transcription switches on when glucose is absent and lactose is present, and off otherwise. Now, let us suppose that we bypass the presence or absence of lactose some way or another to render the repressor either permanently effective or permanently ineffective. We expect that, by making the repressor ineffective, the operon stays on more often than under normal conditions. On the other hand, if the repressor is made effective, we expect the operon to remain off irrespective of the possible glucose deficiency. None of the previous are cases of differential expression. Actually, *Lac* expression is never reaching values that were impossible to take had the repressor’s action not being manipulated externally and independently from the normal environmental cues. Conversely, both are cases of non-differential dysregulations. Even so, only the latter case is classificatory, since the former is tuning the on/off frequencies without quenching the accessible expression levels. The *Lac* example cannot capture the full complexity of the regulatory machinery acting on a Human gene. However, the case of *Lac* is suitable to demonstrate how non-differential dysregulations may emerge from alterations in the behavior of regulators.

The nature of gene dysregulation resides in the lack of normal control of the gene expression, whether it produces aberrant expression values or not. A differentially expressed gene is thus dysregulated because it would not have assumed its actual expression level had it been normally regulated. In contrast, a non-differentially dysregulated gene is indeed dysregulated since it cannot be normally regulated toward certain expression values as a result of an event that would otherwise trigger such behavior. The classificatory cases of non-differential dysregulations correspond to genes that are expressed within normal bounds but are incapacitated to take expression values that are normally available. The latter sets apart from the other plausible case in which non-differential dysregulations amounts to a rearrangement of the gene expression distribution, while maintaining the normal range.

### Homeostasis and cancer

Some tumors are within normal expression bounds even for genes that can be differentially expressed. In fact, 36% of the prostate tumors span the entire range of normal expression for *CTF1* (see Figure 1). In our view, these are not cases of dysregulation since the modification of the relevant expression distribution could be explained by the expression shift in the rest of the tumors. Nonetheless, the 36% of prostate tumors exhibiting a normal regulation of *CTF1* do express differentially at least another member of the panel to which *CTF1* belongs (namely, that with ‘only tumor below’).

As expected, what determines homeostasis is not the normal regulation of individual genes but a collective pattern of normal regulation across a group of them that we called a normal concept. Normal concepts are quite large, involving over a fifth of the genome and as much as half of it in some cases (see Table I). Still, individual genes that fall within normal concepts are strongly interconnected so as that a dysregulation in one of them is always co-instantiated with dysregulations in many others. In fact, the least dysregulated tumor sample in PRAD features 216 dysregulations within the normal concept (i.e., ~1% of the normal concept). Such interdependence allows us to probe the homeostatic state by means of a gene panel which is considerably (i.e., three orders of magnitude) smaller than the collective.

Any dysregulation within the normal concept entails a tumor. To wit, it is essential that normal concept members allows for a dysregulation-based classification which is free from false tumors. In order to exclude genes with low tumor sensitivity, we also demand that normal-concept genes pass a Fisher’s test with a confidence level of 99%. In PRAD, the resulting genes are dysregulated in at least 9% (45/499) of the tumor samples. It is rather unlikely that noise or error behind the data (inherent to biological processes or measurements) could compromise such a robust membership to the normal concept.

Genes within the normal concept lack false tumors, but they do not necessarily classify *all* tumors. However, we may assemble a panel of normal-concept genes that jointly cover all tumor instances. Starting from the most frequently dysregulated gene of the latter kind, such panel is built by incorporating sequentially genes (of the same kind) that are most frequently dysregulated within the yet undetected tumors. This process runs until the ensuing classification is perfect. When the gene to be included in the panel is not univocally determined by the selection criterion, we prioritize genes that are more likely to be dysregulated in general, therefore, enforcing co-dysregulation redundancies. We expect this strategy to yield panels that are less prone to errors in potential diagnostic applications.

Not only is it possible for a prostate tumor to express a gene like *CTF1* within normal bounds, but it is also possible for a normal prostate to express genes (such as *HJURP*) within its tumor scope. Consistently with a case of non-differential dysregulation, 31% of the normal samples express *HJURP* as tumors do (see Figure 1). For all we know, such might be instances of non-differential dysregulations just as much as the ones associated with tumor samples. Even if they are indeed so, the aforementioned 31% of normal samples is normally regulated in at least another member of the panel to which *HJURP* belongs (namely, that with ‘only normal below’).

Like homeostasis, cancer is an ordered state constituted by the joint operation of many genes. Common to all cases of cancer is not the dysregulation of a single gene but a collective pattern of dysregulations across a group of them that we called a tumor concept. In average, tumor concepts are an order of magnitude smaller than normal concepts. On the other hand, genes that fall within tumor concepts are not as strongly interconnected as their normal-concept counterparts. This becomes clear from the fact that the most dysregulated normal samples are only 7 genes (i.e., ~0.3% of the tumor concept) apart from the cancer state. However, the tumor concept is interdependent enough to allow a faultless classification of the state by means of a gene panel, in this case two orders of magnitude smaller than the collective.

In analogy with the way we build the normal concept, any gene is a member of the tumor concept so long as its classification is free from false normal samples and it passes the Fisher’s test with a confidence level of 99%. The latter yields a lower bound of normal regulation instances which is generally too meager. For PRAD, we obtain tumor-concept genes that are normally regulated in at least 2 (4%) of the normal samples. Therefore, tumor concept membership should be less robust than that of the normal concept.

Even if free from false normal classifications, a gene from the tumor concept may not guarantee a thorough detection of normal samples. We overcome this limitation by assembling tumor-concept genes into a panel that collectively detect all normal cases. The strategy here is to select (out of the tumor concept) the gene that instantiate more frequently a normal regulation within the yet unclassified normal samples, until the classification is perfect. If ambiguities emerge, we prioritize genes that instantiate as much of the normal regulations as possible. This sort of co-expression redundancy is not only aimed at compiling a robust diagnostic panel, but also at suggesting optimal candidates to be intervened upon in preclinical development.

### Carcinogenesis and cancer gene therapy

In principle, the normal concept must contain genes involved in carcinogenesis, since these are the ones that lead to cancer upon experiencing a dysregulation. However, we cannot ensure that we are accounting for the relevant drivers due to the strong statistical significance constraints we set up for the discovery of member of the normal concept. Indeed, recall that driver mutations can be rare, and that the minimum number of tumor samples a normal-concept gene is allowed to classify correctly is about 10% of the total amount of tumors within the data.

Nevertheless, the possibility of finding drivers is by no means excluded. In fact, all normal concept genes may well be understood as causally connected to cancer that are more or less efficacious as, and/or are more or less prone to be *initiators* of carcinogenesis. In other words, it may still be the case that the malignant state develops from an intervention that dysregulates any of the genes belonging to the normal concept, regardless of their dysregulation frequency in observational data (such as TCGA’s).

Carcinogenesis have no analogous notion in the case of the tumor concept. Despite the fact that it is sufficient to detect a single, normally regulated gene from a tumor concept to identify a normal sample, the homeostatic state does not grow naturally out of a tumor given appropriate driver regulations. Yet, tumor-concept genes may be understood as causally connected to homeostasis, in the same way as normal-concept genes are causally connected to cancer. Indeed, our observational data supports the hypothesis that sustaining a normal regulatory behavior in a tumor-concept gene within a malignant cell may produce an unviable cell or revert it to a non-tumorous state. This hypothesis is consistent with the strategy envisaged by Kauffman (1971) and developed by Huang and Kauffman (2013) to treat cancer.

### Concluding remarks and outlook

We have shown that it is possible to set up a combinatorial gene panel acting as a perfect biomarker for cancer. That is, monitoring a gene expression profile for such panel members we can correctly ascribe the sample a normal or a tumorous status. In particular, it is possible to classify a sample as tumorous from a single over-expression. However, the latter is but a case among eight sorts of panels, all of which are highly sensitive and specific.

We study 12 cancer types from *The Cancer Genome Atlas* database, among which we include the most common in the world. Five to eight perfect panels are provided per cancer type, depending on the case. In addition, we suggest two other panels combining classifier genes without false tumors, or without false normal samples. A full inventory of such panels can be found in the Supplementary Information, along with a detailed account of our methodology and the robustness of our findings. We expect that this information can be straightforwardly translated into flexible and effective diagnostic panels.

Our gene discovery stems from a view of gene dysregulation beyond the paradigm of differential expression. We start from granting that a dysregulated gene may even feature normal expression values. Indeed, as a result of certain kinds of dysregulation a gene may be *unable to regulate* upon environmental cues towards a given sector within the normal expression window. Therefore, dysregulation is not only constituted by an aberration in the gene expression, but also by a possible limitation in its scope. We dub the latter as *non-differential* dysregulation, in contrast with the term (i.e., *differential* expression) typically used to describe the former.

Furthermore, we stress that differential expression does not only associate with gene expression levels: that are either under or over a reference value, that are demarcated by fixed thresholds, and that are present in most abnormal samples. Take the following counterexample to the first point. A circadian clock gene may be differentially expressed when its expression oscillation stops, even if the halting just yields reference expression values. Now, we summarize an argument against using fixed thresholds. Since gene products operate differently within the organism, gene regulation strategies should reflect such differences somehow, e.g., by customizing normal expression thresholds.

Challenges to the majority constraint come from realizing that coarse-grained phenotypic differences between normal and abnormal tissues are underpinned by multivariate alterations in overall expression profiles. In particular, we note that the abnormal character of a particular sample may not be represented in every gene, and that, for any given gene, an abnormal sample may display an altered expression where another one does not. In fact, two independent abnormal samples may not share the genetic origin of their abnormality. More and more observed instances of altered expression for some particular gene are thus to be taken as better and better evidence of the differential expression taking place for that gene, not as providing the grounds to rule it out. Moreover, taking the majority constraint may overshoot for the required statistical significance. Actually, in the TCGA cohort for PRAD, a 10% of the tumor samples with expression values going beyond a certain threshold is indicative enough of the differential expression taking place for the gene in question.

The proposed view of gene dysregulation allows us to establish broad properties of the gene regulatory network in its homeostatic and cancer states that are consistent with known cancer epidemiology. In particular, homeostasis is characterized by a much greater collection of genes than cancer, thus making carcinogenesis (a departure from homeostasis) substantially more likely than the reverse. Remarkably, identifying minimal assemblies of genes that define cancer might be a first step towards a possible therapy. For instance, it could be that an intervention on non-differentially dysregulated genes either induces a reversion of a malignant cell to its previous normal status, or simply kills it. However, better grounds for such a claim requires additional evidence of causal connections between the involved genes and cancer.

## Acknowledgements

The authors acknowledge support by the Agency of Nuclear Energy and Advanced Technologies (AENTA) of Cuba and the Office of External Activities from the Abdus Salam International Center for Theoretical Physics (ICTP). We are in debt with the members of the ad-hoc Theoretical and Computational Biology group, especially Y. Perera, J. Fernández de Cossío, D. Vázquez, J. Mato, K. Alfaro, J.P. Gómez, J. Nieves and R. Herrero for participating in the preliminary discussions leading to this paper. G.G. is also grateful to L. Azor for her careful reading and most useful suggestions.

## Supplementary Information

### A. Inventory of perfect gene panels

Hereafter, we list the perfect gene panels obtained for the cancer types under study. Panel naming is suggestive of the classificatory feature of their members (see main text). We use Entrez symbols to identify single genes, whenever available. Alternatively, we employ the Ensembl code. For each panel, genes are itemized in descending order of the number of type II errors that one commits when performing a classification based on their expression level. Panels are accompanied by a set of reference expression values and another set of expression thresholds or intervals, which are essential for our classification scheme. Each element of the latter sets stands in a one-to-one correspondence with a panel member in their order of appearance. Reference expression is given in Fragments per Kilobase of transcript per Million mapped reads (FPKM). We use the geometric mean of expression values for normal samples to estimate the reference level. Expression thresholds are given in the binary logarithm of the fold change with respect to the corresponding reference. We denote an interval by bracketed pair of down and up thresholds. Note that numerical precision in reference and threshold values is arbitrary. For ‘only x in zone’ panels, we indicate the character of the classifier gene by appending ‘(a)’, ‘(b)’, ‘(o)’ or ‘(i)’ to the corresponding threshold or interval. We use ‘(a)’ or ‘(b)’ whenever the exclusive expression levels for type x samples lie above (a) or below (b) the threshold it attaches to. Analogously, we use ‘(o)’ or ‘(i)’ whenever the exclusive expression levels are outside (o) or inside (i) the associated interval.

#### A1. Base method (see Section B)

##### Cancer type

LIHC (50 normal samples, 374 tumor samples)

<u>Panel</u> ‘Only tumor above’ (3 genes)

*GABRD*, *SEPTIN7P2* and *TOMM40L*

Reference (FPKM) = {0.153629, 0.577502, 2.08713}

Threshold (log fold change) = {0.949913, 0.752122, 0.859442}

<u>Panel</u> ‘Only tumor below’ (5 genes)

*ANGPTL6, LCAT, UGT2B4, LINC02027* and *MIP*

Reference (FPKM) = {7.31164, 65.2771, 235.995, 5.40384, 0.302426}

Threshold (log fold change) = {−0.939282, −1.16712, −0.836349, −1.19357, −0.605779}

<u>Panel</u> ‘Only normal below’ (5 genes)

*LRRC14*, *MSTO1*, *AURKA*, *APLN* and *FLAD1*

Reference (FPKM) = {1.14623, 0.707592, 0.661075, 0.193745, 4.36832}

Threshold (log fold change) = {0.171096, 0.174825, −0.0568524, −0.112109, 0.0279561}

<u>Panel</u> ‘Only tumor outside’ (4 genes)

*CLDN10*, *HTRA3*, *ALKAL2* and *GCGR*

Reference (FPKM) = {0.97041, 0.676429, 0.573174, 18.7175}

Thresholds (log fold change) = {{−1.50682, 2.1725}, {−1.32299, 1.22535}, {−1.23263, 1.25317}, {−1.20764, 0.710398}}

<u>Panel</u> ‘Only tumor inside’ (6 genes)

*CAVIN4*, *TRPC6*, *AC020765.2*, *COL7A1*, *CCT4* and *CARM1*

Reference (FPKM) = {0.124711, 0.123214, 0.327866, 0.194052, 18.9608, 3.32131}

Thresholds (log fold change) = {{0.32464, 2.05162}, {0.306089, 1.22853}, {0.831366, 1.77933}, {0.568283, 3.57636}, {0.425489, 1.16832}, {0.469648, 1.236}}

<u>Panel</u> ‘Only tumor in zone’ (3 genes)

*GABRD*, *ANGPTL6* and *CEP131*

Reference (FPKM) = {0.153629, 7.31164, 0.764031}

Threshold (log fold change) = {0.949913 (a), −0.939282 (b), 0.821381 (a)}

<u>Panel</u> ‘Only normal in zone’ (5 genes)

*LRRC14*, *MSTO1*, *MARCO*, *VPS45* and *APLN*

Reference (FPKM) = {1.14623, 0.707592, 26.2951, 1.43707, 0.193745}

Threshold (log fold change) = {0.171096 (b), 0.174825 (b), 1.04941 (a), 0.113743 (b), −0.112109 (b)}

##### Cancer type

BRCA (112 normal samples, 1096 tumor samples)

<u>Panel</u> ‘Only tumor above’ (6 genes)

*MMP11, FLAD1, SLC7A8, CCN4, PTTG1IP* and *RRP1B*

Reference (FPKM) = {0.767723, 7.07262, 10.9601, 0.491245, 75.9409, 8.6189}

Threshold (log fold change) = {3.62283, 0.54214, 1.25674, 2.24687, 0.586878, 0.559971}

<u>Panel</u> ‘Only normal above’ (10 genes)

*ARHGAP20, VEGFD, PAMR1, AC084759.3, LEPR, SLC2A4, DST, CBX7, ELOVL7* and *FAM236D*

Reference (FPKM) = {2.74639, 13.1902, 16.771, 0.353627, 6.54067, 4.43987, 23.0258, 10.7712, 3.26935, 0.122595}

Threshold (log fold change) = {−0.176049, 0.255401, 0.451629, 1.15917, 0.303739, 0.353871, 0.874836, 0.594727, 2.92414, 1.53963}

<u>Panel</u> ‘Only tumor below’ (6 genes)

*SPRY2*, *CA4*, *TMEM220-AS1*, *PGM5P3-AS1*, *FP325317.1* and *GABARAPL1*

Reference (FPKM) = {21.6498, 6.21766, 1.03597, 0.299039, 1.4278, 27.4065}

Threshold (log fold change) = {−0.904882, −2.91738, −0.812144, −1.13653, −3.4059, −0.916223}

<u>Panel</u> ‘Only tumor outside’ (8 genes)

*AGTR1, SMOC1, KLHDC9, GXYLT2, GSTA4, ULBP2, MAPT* and *BOLA3-AS1*

Reference (FPKM) = {5.53076, 2.68403, 2.16846, 3.371, 11.2603, 0.385113, 4.59755, \ 2.57972}

Thresholds (log fold change) = {{−1.98996, 2.21559}, {−1.88025, 1.93279}, {−1.67776, 1.29533}, {−1.36319, 1.42113}, {−0.7729, 0.754477}, {−1.24327, 1.26308}, {−2.33635, 1.64319}, {−1.17824, 0.688438}}

<u>Panel</u> ‘Only tumor inside’ (19 genes)

*MUCL1*, *SDK2*, *AL353693.1*, *KRT17*, *SDS*, *H4C8*, *KRT14*, *RSPO1*, *PI15*, *SOD3*, *ELF5*, *HSPB6*, *MMP7, AC010307.4, H4C12, NKAIN1, AC091057.1, LRP2* and *MMP13*

Reference (FPKM) = {105.787, 1.30366, 1.01618, 97.5077, 0.62832, 1.88858, 144.502, \ 1.42586, 10.5847, 32.3513, 9.66864, 69.0263, 21.9762, 0.111588, \ 0.251048, 0.561635, 0.293695, 3.75877, 0.171532}

Thresholds (log fold change) = {{−7.28529, −3.81449}, {−2.44265, −1.17139}, {−2.9087, −1.71572}, {−8.31787, −4.84987}, {2.41369, 3.17379}, {2.59179, 3.9546}, {−4.20293, −2.16278}, {−3.31743, −2.38234},

{−3.81738, −2.3662}, {−2.27724, −1.63909}, {−5.61394, −3.97991}, {−3.34317, −2.46666}, {−3.47107, −2.1004}, {0.946803, 1.71598}, {1.15449, 1.71199}, {2.35936, 3.19706},

{0.879363, 1.4135}, {−5.1978, −4.46481}, {2.06962, 3.10462}}

<u>Panel</u> ‘Only tumor in zone’ (4 genes)

*SPRY2*, *CILP2*, *ITIH5* and *BNIP3P11*

Reference (FPKM) = {21.6498, 0.435807, 15.6612, 0.983961}

Threshold (log fold change) = {−0.904882 (b), 1.65373 (a), −1.99993 (b), 1.62433 (a)}

<u>Panel</u> ‘Only normal in zone’ (10 genes)

*ARHGAP20, VEGFD, PAMR1, AC084759.3, C1QTNF9, DST, P4HA3, LINC01594, ZNF300P1* and *FAM236D*

Reference (FPKM) = {2.74639, 13.1902, 16.771, 0.353627, 0.997502, 23.0258, 0.467047, 0.172191, 1.57727, 0.122595}

Threshold (log fold change) = {−0.176049 (a), 0.255401 (a), 0.451629 (a), 1.15917 (a), 0.539301 (a), 0.874836 (a), −1.52681 (b), 5.17299 (a), 2.29992 (a), 1.53963 (a)}

##### Cancer type

COAD (41 normal samples, 473 tumor samples)

<u>Panel</u> ‘Only tumor above’ (2 genes)

*KRT80* and *ESM1*

Reference (FPKM) = {0.175384, 0.144671}

Threshold (log fold change) = {1.4392, 1.15576}

<u>Panel</u> ‘Only normal above’ (1 gene)

*SCARA5*

Reference (FPKM) = 19.6351

Threshold (log fold change) = −1.31936

<u>Panel</u> ‘Only tumor below’ (1 gene)

Same as panel with ‘only normal above’.

<u>Panel</u> ‘Only normal below’ (2 genes)

*AJUBA* and *KRT80*

Reference (FPKM) = {0.585832, 0.175384}

Threshold (log fold change) = {0.764301, 0.977028}

<u>Panel</u> ‘Only tumor outside’ (3 genes)

*GABRB3*, *ACSL6* and *MTDH*

Reference (FPKM) = {0.374165, 0.180111, 18.0182}

Thresholds (log fold change) = {{−0.799193, 0.931685}, {−0.457481, 0.536373}, {−0.595102, 0.30768}}

<u>Panel</u> ‘Only tumor inside’ (3 genes)

*SEC14L2, TMEM41A* and *FABP6*

Reference (FPKM) = {0.373755, 2.29925, 0.322822}

Thresholds (log fold change) = {{0.655081, 4.09315}, {0.498709, 2.12734}, {3.43029, 9.5477}}

<u>Panel</u> ‘Only tumor in zone’ (1 gene)

Same as panel with ‘only tumor below’.

<u>Panel</u> ‘Only normal in zone’ (1 gene)

Same as panel with ‘only normal above’.

##### Cancer type

HNSC (44 normal samples, 502 tumor samples)

<u>Panel</u> ‘Only tumor above’ (4 genes)

*CDCA5*, *GPRIN1*, *LINC01633* and *OFCC1*

Reference (FPKM) = {2.46466, 0.847135, 0.105283, 0.103059}

Threshold (log fold change) = {1.31901, 1.63809, 0.472287, 0.217898}

<u>Panel</u> ‘Only normal above’ (8 genes)

*EMP1, PIP, KRTAP13-1, CIDEC, LINC00443, ANXA1, CLEC3B* and *CYP3A4*

Reference (FPKM) = {154.301, 6.93504, 0.232112, 0.857577, 0.208645, 770.135, 10.0374, 0.274655}

Threshold (log fold change) = {0.909014, 5.58461, 1.03298, 1.44621, 0.271165, 1.3106, 1.0025, 2.02738}

<u>Panel</u> ‘Only tumor below’ (5 genes)

*ADIPOQ*, *CYP4B1*, *TPT1*, *GPT2* and *B4GALT1-AS1*

Reference (FPKM) = {1.05582, 9.91754, 470.502, 19.5348, 1.34636}

Threshold (log fold change) = {−3.32081, −2.72798, −0.602725, −1.12478, −0.841166}

<u>Panel</u> ‘Only tumor outside’ (6 genes)

*MSL3P1, THSD7B, RNF144A, RPL39L, SAP18* and *ANAPC5*

Reference (FPKM) = {0.196208, 0.303781, 1.76498, 3.44229, 26.7148, 7.60304}

Thresholds (log fold change) = {{−0.790677, 0.599432}, {−1.19036, 1.8796}, {−1.11365, 1.03681}, {−1.89815, 1.65638}, {−0.369018, 0.738929}, {−0.264216, 0.370855}}

<u>Panel</u> ‘Only tumor inside’ (5 genes)

*BLOC1S3*, *AC109479.1*, *PDZK1*, *KRTAP4-1* and *HOMER3*

Reference (FPKM) = {3.32552, 0.125148, 0.17184, 0.134966, 4.31855}

Thresholds (log fold change) = {{0.713113, 1.68529}, {0.75768, 3.88775}, {0.224232, 5.63674}, {1.26071, 6.37495}, {1.18186, 2.59481}}

<u>Panel</u> ‘Only tumor in zone’ (4 genes)

*CDCA5*, *ATP5F1A*, *OFCC1* and *RFC4*

Reference (FPKM) = {2.46466, 54.5384, 0.103059, 2.66874}

Threshold (log fold change) = {1.31901 (a), −0.554119 (b), 0.217898 (a), 0.825152 (a)}

<u>Panel</u> ‘Only normal in zone’ (6 genes)

*MMP9*, *GPRIN1*, *FHOD1*, *BMP1*, *CIDEC* and *EMP1*

Reference (FPKM) = {2.18071, 0.847135, 3.14689, 2.95999, 0.857577, 154.301}

Threshold (log fold change) = {2.18071 (b), 0.847135 (b), 3.14689 (b), 2.95999 (b), 0.857577 (a), 154.301 (a)}

##### Cancer type

KIRC (74 normal samples, 539 tumor samples)

<u>Panel</u> ‘Only tumor above’ (4 genes)

*DDB2, PARVB, SLC15A4* and *GACAT2*

Reference (FPKM) = {2.72934, 2.73299, 4.8592, 0.166507}

Threshold (log fold change) = {0.890201, 1.02849, 1.12276, 1.29121}

<u>Panel</u> ‘Only normal above’ (3 genes)

*AQP2*, *SOST* and *TSPAN6*

Reference (FPKM) = {529.234, 2.75347, 25.8478}

Threshold (log fold change) = {−2.05659, 0.302995, 0.546245}

<u>Panel</u> ‘Only tumor below’ (3 genes)

*AC104237.2*, *PAQR7* and *LY86-AS1*

Reference (FPKM) = {2.86161, 23.245, 0.193608}

Threshold (log fold change) = {−1.32775, −0.932102, −0.599823}

<u>Panel</u> ‘Only tumor outside’ (4 genes)

*PCDHA11*, *TGFA*, *MACROD2* and *PGBD5*

Reference (FPKM) = {0.436441, 9.83772, 1.1507, 0.76372}

Thresholds (log fold change) = {{−0.94271, 0.982452}, {−1.02951, 1.02527}, {−0.842658, 0.585687}, {−1.01488, 1.07635}}

<u>Panel</u> ‘Only tumor inside’ (7 genes)

*NEIL3*, *BCAN*, *LY6E*, *CENPU*, *LAT2*, *NKAIN1* and *AC073869.1*

Reference (FPKM) = {0.139495, 0.223697, 27.0516, 0.61479, 0.799776, 0.148608, 0.142873}

Thresholds (log fold change) = {{0.578049, 2.22675}, {0.89292, 2.16564}, {1.09067, 2.63392},

{1.06547, 2.39031}, {1.64292, 2.3807}, {0.689439, 2.54531}, {0.85847, 1.5176}}

<u>Panel</u> ‘Only tumor in zone’ (3 genes)

Same as panel with ‘only tumor below’.

<u>Panel</u> ‘Only normal in zone’ (3 genes)

Same as panel with ‘only normal above’.

##### Cancer type

KIRP (32 normal samples, 289 tumor samples)

<u>Panel</u> ‘Only tumor above’ (3 genes)

*HK2*, *AC124798.1* and *NME1*

Reference (FPKM) = {0.89216, 1.33416, 4.97076}

Threshold (log fold change) = {1.55936, 0.760797, 0.652767}

<u>Panel</u> ‘Only normal above’ (1 gene)

*UMOD*

Reference (FPKM) = 1882.42

Threshold (log fold change) = −5.42522

<u>Panel</u> ‘Only tumor below’ (1 gene)

Same as panel with ‘only normal above’.

<u>Panel</u> ‘Only tumor outside’ (3 genes)

*RGS12, AC090204.1* and *PREX2*

Reference (FPKM) = {3.17738, 1.71275, 0.923607}

Thresholds (log fold change) = {{−0.501801, 0.461124}, {−1.03531, 0.835089}, {−0.85281, 1.22689}}

<u>Panel</u> ‘Only tumor inside’ (6 genes)

*TTTY15, SRSF3P1, PRDX4, CHPF, TAP1* and *AC008760.2*

Reference (FPKM) = {0.554073, 0.562639, 17.9703, 29.4517, 9.46457, 0.1918}

Thresholds (log fold change) = {{−2.43075, 0.604354}, {−1.81578, −0.857145}, {0.464816, 1.08214}, {0.981492, 2.31435}, {0.882734, 1.61068}, {0.973484, 3.13367}}

<u>Panel</u> ‘Only tumor in zone’ (1 gene)

Same as panel with ‘only tumor below’.

<u>Panel</u> ‘Only normal in zone’ (1 gene)

Same as panel with ‘only normal above’.

##### Cancer type

LUAD (59 normal samples, 535 tumor samples)

<u>Panel</u> ‘Only tumor above’ (3 genes)

*ALDH18A1*, *TRIM27* and *PYCR1*

Reference (FPKM) = {9.25462, 6.24107, 2.89561}

Threshold (log fold change) = {0.745953, 0.311544, 1.54081}

<u>Panel</u> ‘Only normal above’ (5 genes)

*STX11*, *CALCRL*, *SFTPC*, *FHL1* and *EDNRB*

Reference (FPKM) = {26.2114, 29.2525, 7522.07, 54.2349, 30.3858}

Threshold (log fold change) = {−0.243529, 0.0562003, 0.0695837, 0.056593, 0.10761}

<u>Panel</u> ‘Only tumor below’ (4 genes)

*AGER*, *ST7*, *FABP4* and *GYPE*

Reference (FPKM) = {1003.51, 9.13645, 85.6647, 1.94262}

Threshold (log fold change) = {−2.04955, −0.360178, −2.93279, −1.29594}

<u>Panel</u> ‘Only tumor outside’ (3 genes)

*PODXL2*, *BEX4* and *RAD23B*

Reference (FPKM) = {2.41595, 37.122, 30.3231}

Thresholds (log fold change) = {{−0.824902, 0.986005}, {−0.396451, 0.321376}, {−0.321894, 0.355567}}

<u>Panel</u> ‘Only tumor inside’ (6 genes)

*NLN*, *GLMP*, *TEX19*, *MMP11*, *IGHV5-78* and *FANCD2OS*

Reference (FPKM) = {1.34552, 9.90187, 0.10632, 0.357485, 0.455233, 0.10663}

Thresholds (log fold change) = {{0.747499, 2.08854}, {0.619486, 1.59271}, {0.19985, 2.6173}, {2.80669, 5.05784}, {2.61737, 4.80558}, {0.202047, 0.806609}}

<u>Panel</u> ‘Only tumor in zone’ (2 genes)

*AGER* and *SLC37A4*

Reference (FPKM) = {1003.51, 3.07233}

Threshold (log fold change) = {−2.04955 (b), 0.389303 (a)}

<u>Panel</u> ‘Only normal in zone’ (5 genes)

Same as panel with ‘only normal above’.

##### Cancer type

LUSC (49 normal samples, 502 tumor samples)

<u>Panel</u> ‘Only tumor above’ (2 genes)

*MRGBP* and *F12*

Reference (FPKM) = {4.11491, 0.352769}

Threshold (log fold change) = {0.405015, 0.876742}

<u>Panel</u> ‘Only normal above’ (3 genes)

*EMP2*, *ESAM* and *GKN2*

Reference (FPKM) = {138.996, 58.6668, 26.7761}

Threshold (log fold change) = {−0.767637, −0.306594, −0.48887}

<u>Panel</u> ‘Only tumor below’ (2 genes)

*ADGRF5* and *ADGRD1*

Reference (FPKM) = {71.9583, 5.83605}

Threshold (log fold change) = {−0.630642, −1.06124}

<u>Panel</u> ‘Only normal below’ (3 genes)

*SLC2A1*, *RFC4* and *PPP1R14BP3*

Reference (FPKM) = {3.59379, 1.90837, 8.07671}

Threshold (log fold change) = {0.646135, 0.305688, −0.0675405}

<u>Panel</u> ‘Only tumor outside’ (3 genes)

*CES1*, *AKR1C2* and *CSAD*

Reference (FPKM) = {76.8643, 1.51769, 2.32698}

Thresholds (log fold change) = {{−1.05284, 1.01772}, {−1.22746, 1.41218}, {−1.6687, 1.2517}}

<u>Panel</u> ‘Only tumor inside’ (6 genes)

*AGMAT*, *SLC16A1*, *CNP*, *PTGES3P3*, *PADI3* and *PHF6*

Reference (FPKM) = {0.309999, 2.69623, 8.64472, 1.42205, 0.123148, 2.40495}

Thresholds (log fold change) = {{1.11876, 3.01349}, {1.73048, 3.49603}, {0.373249, 1.05463}, {0.826624, 1.71155}, {0.384183, 4.51551}, {0.601766, 1.33932}}

<u>Panel</u> ‘Only tumor in zone’ (2 genes)

Same as panel with ‘only tumor below’.

<u>Panel</u> ‘Only normal in zone’

*EMP2* and *HDAC2*

Reference (FPKM) = {138.996, 3.98041}

Threshold (log fold change) = {−0.767637 (a), −0.00464922 (b)}

##### Cancer type

PRAD (52 normal samples, 499 tumor samples)

<u>Panel</u> ‘Only tumor above’ (8 genes)

*EPHA10, UCN, TMEM86A, RPL7AP31, FRMPD3, RP11-658F2.8, HOXA10-AS* and *UBXN6*

Reference (FPKM) = {0.391225, 0.571144, 1.57228, 0.16683, 0.343019, 4.09354, 0.975031, 30.2444}

Threshold (log fold change) = {1.3373, 1.33742, 0.761516, 0.883756, 1.27324, 0.607416, 0.831813, 0.420475}

<u>Panel</u> ‘Only normal above’ (14 genes)

*SEPTIN10, KLHL4, CD82, LINC01546, HCAR2, FGD5P1, MYH11, SLC18A2, RNA5SP342, AL161668.4, TES, ASS1P1, TRMT1L* and *AC093536.1*

Reference (FPKM) = {16.8358, 0.30878, 22.5065, 0.255483, 1.7208, 0.163109, 433.746, 1.31571, 0.150767, 0.337145, 29.5951, 0.327261, 4.89661, 0.500265}

Threshold (log fold change) = {0.426986, 0.553361, 0.829505, 1.07419, 2.12047, 0.601488, 1.07166, 2.78262, 2.24141, 1.01562, 0.669396, 1.5279, 0.488484, 1.77176}

<u>Panel</u> ‘Only tumor below’ (15 genes)

*CTF1*, *SH3GLB1*, *C19orf47*, *DLEU1*, *NRSN2-AS1*, *RHOQP3*, *YWHAZP10*, *CRYAB*, *AP2B1P1*, *TGM5*, *DNAJB4*, *CLCA4-AS1*, *CASQ1*, *GBP6* and *DCTN6*

Reference (FPKM) = {9.36788, 16.5119, 3.90623, 0.583145, 3.43836, 0.222637, 1.00048, 42.1054, 0.138725, 0.293681, 7.18043, 0.616539, 1.7037, 0.235882, 12.3231}

Threshold (log fold change) = {−1.09605, −0.419621, −0.364528, −0.568821, −0.544813, −0.729442, −1.1141, −1.6243, −0.383461, −1.2737, −1.26595, −0.751685, −1.97423, −0.940522, −0.414721}

<u>Panel</u> ‘Only tumor outside’ (10 genes)

*PRRT3-AS1*, *AC245100.2*, *NCALD*, *POU6F2*, *C2orf16*, *CPS1*, *SYCE1L*, *EML4*, *AC124947.1* and *AC111182.1*

Reference (FPKM) = {1.54542, 0.176046, 2.45813, 0.12238, 0.373783, 0.310419, 1.62368, 5.48395, 0.427871, 0.12869}

Thresholds (log fold change) = {{−1.27757, 1.37632}, {−0.692421, 0.944042}, {−1.21066, 1.22599}, {−0.266019, 0.866407}, {−0.566091, 0.740341}, {−0.862699, 1.88542}, {−1.37765, 1.49611}, {−0.924394, 0.95464}, {−1.10092, 0.827269}, {−0.294208, 0.707256}}

<u>Panel</u> ‘Only tumor inside’ (8 genes)

*SNHG3, SAMD5, DLX2, DUS1L, NETO2, TMEM25, AC106772.2* and *AC073548.1*

Reference (FPKM) = {2.54932, 1.29739, 0.150705, 14.3955, 0.393405, 7.06784, 0.110755, 0.539203}

Thresholds (log fold change) = {{1.06334, 2.52785}, {1.0579, 3.74196}, {0.694075, 2.5946}, {0.921421, 2.26158}, {0.581784, 1.85239}, {0.410563, 1.20698}, {0.581043, 2.29143}, {0.852799, 2.22106}}

<u>Panel</u> ‘Only tumor in zone’ (7 genes)

*EPHA10*, *UCN*, *DLX2*, *FAM86B3P*, *COL10A1*, *SDCBP2-AS1* and *AC099518.1*

Reference (FPKM) = {0.391225, 0.571144, 0.150705, 0.520009, 0.242829, 1.01677, 0.45783}

Threshold (log fold change) = {1.3373 (a), 1.33742 (a), {0.694075, 2.5946} (i), {−0.677684, 0.724937} (o), {2.54639,4.27342} (i), {−1.04495, 0.687429} (o), {0.956875, 2.20259} (i)}

<u>Panel</u> ‘Only normal in zone’ (11 genes)

*HJURP*, *SEPTIN10*, *AL512363.1*, *CA14*, *FGD5P1*, *SEPTIN5*, *TES*, *SLC18A2*, *SNORD104*, *TBC1D7* and *RNA5SP342*

Reference (FPKM) = {0.277751, 16.8358, 0.446229, 2.11777, 0.163109, 2.5737, 29.5951, 1.31571, 4.33524, 1.466, 0.150767}

Threshold (log fold change) = {−0.610867 (b), 0.426986 (a), 0.925758 (a), 0.8031 (a), 0.601488 (a), −1.20939 (b), 0.669396 (a), 2.78262 (a), −2.30014 (b), −0.592224 (b), 2.24141 (a)}

##### Cancer type

STAD (32 normal samples, 375 tumor samples)

<u>Panel</u> ‘Only tumor above’ (3 genes)

*ESM1, AMH* and *ZNF761*

Reference (FPKM) = {0.183189, 0.250209, 1.54561}

Threshold (log fold change) = {0.946779, 1.09099, 0.711567}

<u>Panel</u> ‘Only normal above’ (5 genes)

*IGHV3OR16-13, DPT, MALL, MT-TY* and *MYZAP*

Reference (FPKM) = {4.59954, 35.5918, 4.16596, 52.7855, 3.04359}

Threshold (log fold change) = {3.868, 1.27223, 3.21166, 2.67099, 0.691261}

<u>Panel</u> ‘Only tumor below’ (5 genes)

*PLP1*, *ALDOC*, *NSG1*, *FZD9* and *KRT222*

Reference (FPKM) = {2.09738, 11.5507, 2.12832, 0.377841, 0.250353}

Threshold (log fold change) = {−2.46448, −1.15265, −1.44638, −1.12761, −0.605064}

<u>Panel</u> ‘Only normal below’ (5 genes)

*CENPL*, *COL10A1*, *XPO5*, *ACAN* and *HSPD1P1*

Reference (FPKM) = {0.802347, 0.166095, 3.18288, 0.165321, 0.34746}

Threshold (log fold change) = {−0.121296, −0.261213, −0.0716832, −0.328354, −0.545085}

<u>Panel</u> ‘Only tumor outside’ (6 genes)

*NUP62CL, AC009237.3, FBXO2, GREB1, MNS1* and *RANBP17*

Reference (FPKM) = {0.466601, 0.414421, 3.1556, 0.283874, 0.92088, 0.215234}

Thresholds (log fold change) = {{−1.34013, 1.18665}, {−0.971996, 0.96658}, {−1.17493, 2.43909}, {−1.12622, 1.13218}, {−1.27687, 0.818611}, {−0.803521, 0.81078}}

<u>Panel</u> ‘Only tumor inside’ (5 genes)

*ZSWIM4, ESPL1, EFNA3, AC008808.2* and *PKP1*

Reference (FPKM) = {2.69178, 1.16598, 1.0856, 1.4811, 0.526516}

Thresholds (log fold change) = {{0.44067, 2.0316}, {0.597465, 2.66237}, {0.339062, 3.14304}, {−3.59924, −1.03791}, {−1.40625, 4.8887}}

<u>Panel</u> ‘Only tumor in zone’ (3 genes)

Same as panel with ‘only tumor above’.

<u>Panel</u> ‘Only normal in zone’ (4 genes)

*CENPL, JCHAIN, BID* and *ACY1*

Reference (FPKM) = {0.802347, 333.792, 3.12532, 1.76071}

Threshold (log fold change) = {−0.121296 (b), 3.88943 (a), −0.547937 (b), {−1.9371, 2.43198} (o)}

##### Cancer type

THCA (58 normal samples, 510 tumor samples)

<u>Panel</u> ‘Only tumor above’ (5 genes)

*METTL7B*, *GJC1*, *TYW1*, *UNC5B-AS1* and *FHOD1*

Reference (FPKM) = {2.09007, 0.642125, 5.65877, 0.147837, 4.68915}

Threshold (log fold change) = {1.70463, 0.868901, 0.363749, 1.00537, 0.736125}

<u>Panel</u> ‘Only normal above’ (11 genes)

*AC109326.1*, *AL034374.1*, *PIK3C2G*, *BCL2L11*, *LINC01589*, *RIF1*, *AC105105.1*, *GPM6A*, *UGT2B11, AC239804.2* and *PDXDC2P*

Reference (FPKM) = {2.20965, 3.68123, 0.121792, 11.0026, 0.262265, 3.7348, 0.106732, \ 3.80712, 0.566833, 0.140631, 0.150152}

Threshold (log fold change) = {0.385581, 0.273349, 0.28758, 0.321295, 0.87205, 0.619417, 0.377582, 0.651311, 0.733642, 1.21499, 0.418387}

<u>Panel</u> ‘Only tumor below’ (6 genes)

*TFF3*, *GBA3, SLC6A16, TRMO, AC010834.2* and *AC131206.1*

Reference (FPKM) = {656.882, 0.370054, 0.76704, 5.66109, 0.210822, 0.215678}

Threshold (log fold change) = {−2.10598, −0.811088, −0.828981, −0.396442, −0.695244, −0.906661}

<u>Panel</u> ‘Only tumor outside’ (5 genes)

*IGF2BP2*, *CLU*, *ORMDL3*, *GLDN* and *MXRA8*

Reference (FPKM) = {3.3912, 382.801, 35.5455, 0.209372, 15.5592}

Threshold (log fold change) = {{−1.45196, 1.19003}, {−0.87064, 0.778572}, {−0.449449, 0.426211}, {−0.797159, 1.01833}, {−1.58551, 0.880012}}

<u>Panel</u> ‘Only tumor inside’ (5 genes)

*SERINC2, HAPLN1, OIT3, GABRD* and *PTP4A3*

Reference (FPKM) = {12.5867, 0.113916, 0.173946, 0.434782, 4.3927}

Threshold (log fold change) = {{0.92633, 2.62982}, {0.635725, 3.58676}, {1.34265, 3.63698}, {2.45567, 4.05592}, {1.34833, 2.66843}}

<u>Panel</u> ‘Only tumor in zone’ (4 genes)

*METTL7B, MFAP2, AC005083.1* and *FHOD1*

Reference (FPKM) = {2.09007, 1.3338, 3.22342, 4.68915}

Threshold (log fold change) = {1.70463 (a), {−1.59529, 1.32259} (o), {−1.25186, 1.2645} (o), 0.736125 (a)}

<u>Panel</u> ‘Only normal in zone’ (9 genes)

*AC109326.1*, *GPRIN1*, *GRIA1*, *GLI1*, *BCL2L11*, *TNFRSF11B*, *AP003084.1*, *OTUD7B* and *AC239804.2*

Reference (FPKM) = {2.20965, 0.215338, 0.15403, 0.599669, 11.0026, 56.8365, 0.91206, 3.58097, 0.140631}

Threshold (log fold change) = {0.385581 (a), −0.392671 (b), 0.215693 (a), 0.690303 (a), 0.321295 (a), 0.952059 (a), 0.651768 (a), {−1.465, 1.34669} (o), 1.21499 (a)}

##### Cancer type

UCEC (23 normal samples, 552 tumor samples)

<u>Panel</u> ‘Only tumor above’ (1 gene)

*TBC1D7*

Reference (FPKM) = 0.832865

Threshold (log fold change) = 0.577791

<u>Panel</u> ‘Only normal above’ (1 gene)

*PLSCR4*

Reference (FPKM) = 16.3218

Threshold (log fold change) = −0.971837

<u>Panel</u> ‘Only tumor below’ (1 gene)

Same as panel with ‘only normal above’.

<u>Panel</u> ‘Only normal below’ (1 gene)

Same as panel with ‘only tumor above’.

<u>Panel</u> ‘Only tumor outside’ (3 genes)

*RNF10, ERO1B* and *PGK1*

Reference (FPKM) = {28.5136, 2.76261, 38.7264}

Thresholds (log fold change) = {{−0.113732, 0.1091}, {−0.585613, 0.507264}, {−0.5956, 0.257718}}

<u>Panel</u> ‘Only tumor inside’ (4 genes)

*HOXD10*, *MARCKSL1*, *RIPK2* and *SCD*

Reference (FPKM) = {2.98662, 93.5577, 4.6923, 4.59202}

Thresholds (log fold change) = {{−1.56749, 2.20744}, {0.70168, 2.50881}, {0.753443, 2.16763}, {1.8648, 3.47497}}

<u>Panel</u> ‘Only tumor in zone’ (1 gene)

Same as panel with ‘only tumor below’.

<u>Panel</u> ‘Only normal in zone’ (1 gene)

Same as panel with ‘only normal below’.

#### A2. Stringent method (see Section B: *Constraining the search*)

##### Cancer type

LIHC (50 normal samples, 374 tumor samples)

<u>Panel</u> ‘Only tumor above’ (3 genes)

*TOMM40L*, *NOTCH3* and *SEMA3F*

Reference (FPKM) = {2.08713, 0.772191, 1.2413}

Threshold (log fold change) = {0.859442, 1.63185, 0.62631}

<u>Panel</u> ‘Only tumor below’ (5 genes)

*LCAT*, *ANGPTL6*, *CYP2C8*, *DCAF11* and *C8B*

Reference (FPKM) = {65.2771, 7.31164, 420.379, 25.3617, 208.861}

Threshold (log fold change) = {−1.16712, −0.939282, −1.34293, −0.626924, −0.477411}

<u>Panel</u> ‘Only tumor outside’ (4 genes)

*MANEAL*, *HIBADH*, *ARMCX3* and *UBE2H*

Reference (FPKM) = {2.12454, 47.379, 4.58176, 9.59768}

Thresholds (log fold change) = {{−0.824129, 0.775906}, {−0.443783, 0.504112}, {−1.21458, 0.813262}, {−0.641592, 0.553812}}

<u>Panel</u> ‘Only tumor inside’ (8 genes)

*RNF187, LAMTOR4, THAP2, SLC51B, SOWAHA, PPM1M, NEDD9* and *ADA*

Reference (FPKM) = {18.9788, 13.8208, 0.416883, 0.659096, 0.74941, 1.54713, 1.98517, 0.93445}

Thresholds (log fold change) = {{0.491761, 1.61893}, {0.832461, 1.92626},

{0.527624, 1.74859}, {1.63761, 4.15464}, {1.19357, 2.43452}, {1.08155, 2.08608}, {1.44413, 2.46249}, {1.05832, 2.18588}}

<u>Panel</u> ‘Only tumor in zone’ (3 genes)

Same as panel with ‘only tumor above’.

<u>Panel</u> ‘Only normal in zone’ (5 genes)

*LRRC14, MSTO1, MARCO, VPS45* and *CYP2C8*

Reference (FPKM) = {1.14623, 0.707592, 26.2951, 1.43707, 420.379}

Threshold (log fold change) = {0.171096 (b), 0.174825 (b), 1.04941 (a), 0.113743 (b), 0.496438 (a)}

##### Cancer type

BRCA (112 normal samples, 1096 tumor samples)

<u>Panel</u> ‘Only tumor above’ (6 genes)

*MMP11, FLAD1, SLC7A8, CCN4, PTTG1IP* and *RRP1B*

Reference (FPKM) = {0.767723, 7.07262, 10.9601, 0.491245, 75.9409, 8.6189}

Threshold (log fold change) = {3.62283, 0.54214, 1.25674, 2.24687, 0.586878, 0.559971}

<u>Panel</u> ‘Only tumor below’ (7 genes)

*SPRY2, GABARAPL1, C22orf39, CD36, EIF2B3, MRAS* and *ITIH5*

Reference (FPKM) = {21.6498, 27.4065, 5.36262, 85.0498, 9.96411, 21.6607, 15.6612}

Threshold (log fold change) = {−0.904882, −0.916223, −0.61716, −3.40855, −0.515781, −1.59663, −1.99993}

<u>Panel</u> ‘Only tumor outside’ (9 genes)

*MAOB*, *SCCPDH*, *GSTA4*, *BMERB1*, *NMNAT3*, *BCL2*, *ASB16-AS1*, *FRMD3* and *ADAM9*

Reference (FPKM) = {60.0867, 22.1882, 11.2603, 7.58791, 3.61775, 14.2168, 3.38053, 2.99692, 14.2106}

Thresholds (log fold change) = {{−1.61654, 1.2328}, {−1.2423, 0.858307}, {−0.7729, 0.754477}, {−1.27483, 0.782717}, {−0.712546, 0.627419}, {−1.35192, 1.01714}, {−0.931551, 0.72679}, {−1.42067, 1.29465}, {−1.91126, 0.915658}}

<u>Panel</u> ‘Only tumor in zone’ (4 genes)

*SPRY2*, *CILP2*, *CD34* and *ELMO2*

Reference (FPKM) = {21.6498, 0.435807, 27.1067, 7.72436}

Threshold (log fold change) = {−0.904882 (b), 1.65373 (a), −1.59334 (b), {−0.65156, 0.470227} (o)}

##### Cancer type

COAD (41 normal samples, 473 tumor samples)

<u>Panel</u> ‘Only tumor above’ (2 genes)

*PVT1* and *ETV4*

Reference (FPKM) = {0.470726, 0.792852}

Threshold (log fold change) = {0.758725, 1.67762}

<u>Panel</u> ‘Only normal above’ (1 gene)

*SCARA5*

Reference (FPKM) = 19.6351

Threshold (log fold change) = −1.31936

<u>Panel</u> ‘Only tumor below’ (2 genes)

*PLPP1* and *METTL7A*

Reference (FPKM) = {40.5178, 56.1139}

Threshold (log fold change) = {−0.648131, −0.79878}

<u>Panel</u> ‘Only normal below’ (5 genes)

*SLC6A6, DUSP14, AJUBA, CDC25B* and *ENC1*

Reference (FPKM) = {1.90672, 1.70866, 0.585832, 7.94452, 6.74848}

Threshold (log fold change) = {0.118954, −0.0874422, 0.764301, 0.0493737, 0.0359625}

<u>Panel</u> ‘Only tumor outside’ (4 genes)

*SF3A2, PPP2R2A, PIGX* and *SUGT1*

Reference (FPKM) = {18.5175, 5.77076, 4.44438, 3.03511}

Thresholds (log fold change) = {{−0.64547, 0.329928}, {−0.753017, 0.32839}, {−0.377202, 0.28793}, {−0.265226, 0.348177}}

<u>Panel</u> ‘Only tumor inside’ (3 genes)

*SEC14L2, TMEM41A* and *FABP6*

Reference (FPKM) = {0.373755, 2.29925, 0.322822}

Thresholds (log fold change) = {{0.655081, 4.09315}, {0.498709, 2.12734}, {3.43029, 9.5477}}

<u>Panel</u> ‘Only tumor in zone’ (2 genes)

Same as panel with ‘only tumor above’.

<u>Panel</u> ‘Only normal in zone’ (1 gene)

Same as panel with ‘only normal above’.

##### Cancer type

HNSC (44 normal samples, 502 tumor samples)

<u>Panel</u> ‘Only tumor above’ (4 genes)

*CDCA5*, *GPRIN1*, *SLC38A7* and *NMRAL2P*

Reference (FPKM) = {2.46466, 0.847135, 2.08616, 0.415419}

Threshold (log fold change) = {1.31901, 1.63809, 0.752531, 2.11205}

<u>Panel</u> ‘Only normal above’ (8 genes)

*EMP1, PIP, CIDEC, CYP3A4, GBP6, KRT13, CLEC3B* and *SERPINB1*

Reference (FPKM) = {154.301, 6.93504, 0.857577, 0.274655, 55.428, 2264., 10.0374, 169.774}

Threshold (log fold change) = {0.909014, 5.58461, 1.44621, 2.02738, 2.58313, 2.88847, 1.0025, 1.39619}

<u>Panel</u> ‘Only tumor below’ (7 genes)

*TPT1, ITM2A, PEBP1, GSN, MAGI1, PCDH1* and *DMAC2L*

Reference (FPKM) = {19.5348, 470.502, 11.757, 144.047, 66.6051, 2.70486, 18.9987, 1.81833}

Threshold (log fold change) = {−1.12478, −0.602725, −1.76573, −0.450407, −1.08436, −1.29483, −1.4952, −0.579562}

<u>Panel</u> ‘Only tumor outside’ (7 genes)

*TRMT12, CTNNAL1, DANCR, ASH2L, ATP5MG, CA11* and *TSR2*

Reference (FPKM) = {3.70353, 6.56481, 14.2095, 6.80895, 36.3973, 3.71942, 22.168}

Thresholds (log fold change) = {{−0.681307, 0.63304}, {−0.942862, 1.07847}, {−1.10101, 0.704067}, {−0.548412, 0.359279}, {−0.475971, 0.880068}, {−1.20365, 1.35953}, {−0.467133, 0.417771}}

<u>Panel</u> ‘Only tumor inside’ (6 genes)

*WDR54, LAMA3, HOMER3, FADS2, C1QTNF6* and *KRT4*

Reference (FPKM) = {1.64298, 4.02691, 4.31855, 1.05067, 0.677837, 804.76}

Thresholds (log fold change) = {{1.59476, 3.67044}, {1.30952, 3.55378}, {1.18186, 2.59481}, {1.5752, 6.40439}, {2.51246, 3.76921}, {−4.07412, 0.634131}}

<u>Panel</u> ‘Only tumor in zone’ (4 genes)

*CDCA5*, *ATP5F1A*, *ZFPM2-AS1* and *EME1*

Reference (FPKM) = {2.46466, 54.5384, 0.144573, 0.408743}

Threshold (log fold change) = {1.31901 (a), −0.554119 (b), {2.04123, 3.71273} (i), 1.07651 (a)}

<u>Panel</u> ‘Only normal in zone’ (6 genes)

*EMP1*, *BMP1*, *PIP*, *CCL14*, *FHOD1* and *COL4A1*

Reference (FPKM) = {154.301, 2.95999, 6.93504, 0.533505, 3.14689, 5.18571}

Threshold (log fold change) = {0.909014 (a), −0.158925 (b), 5.58461 (a), 1.47779 (a), −0.599631 (b), −0.34559 (b)}

##### Cancer type

KIRC (74 normal samples, 539 tumor samples)

<u>Panel</u> ‘Only tumor above’ (4 genes)

*DDB2, PARVB, SLC15A4* and *NADSYN1*

Reference (FPKM) = {2.72934, 2.73299, 4.8592, 2.03386}

Threshold (log fold change) = {0.890201, 1.02849, 1.12276, 0.674794}

<u>Panel</u> ‘Only normal above’ (3 genes)

*AQP2, SOST* and *TSPAN6*

Reference (FPKM) = {529.234, 2.75347, 25.8478}

Threshold (log fold change) = {−2.05659, 0.302995, 0.546245}

<u>Panel</u> ‘Only tumor below’ (3 genes)

*GABARAPL1, FAM104B* and *RASSF8*

Reference (FPKM) = {81.887, 7.11941, 11.2172}

Threshold (log fold change) = {−0.893622, −0.399217, −0.724517}

<u>Panel</u> ‘Only tumor outside’ (5 genes)

*FOXJ3, UBL3, ATP11A, EIF4B* and *LINC02532*

Reference (FPKM) = {8.22586, 25.7548, 7.72559, 66.6365, 9.06415}

Thresholds (log fold change) = {{−0.485845, 0.421358}, {−0.637267, 0.60413},

{−0.861355, 1.22126}, {−0.819592, 0.417683}, {−0.964831, 0.692758}}

<u>Panel</u> ‘Only tumor inside’ (9 genes)

*CENPU, LY6E, BX640514.2, SKA3, BCAN, CD48, OR51E1, RPL10P9* and *AC099850.3*

Reference (FPKM) = {0.61479, 27.0516, 0.302963, 0.243294, 0.223697, 1.20936, 0.403187, 1.87362, 0.543449}

Thresholds (log fold change) = {{1.06547, 2.39031}, {1.09067, 2.63392}, {1.45249, 3.33541}, {1.35927, 2.86844}, {1.42286, 2.16564}, {2.38149, 3.3886}, {1.67234, 2.7621}, {1.29915, 2.8916}, {1.24845, 2.35761}}

<u>Panel</u> ‘Only tumor in zone’ (3 genes)

*DDB2, PAQR7* and *NADSYN1*

Reference (FPKM) = {2.72934, 23.245, 2.03386}

Threshold (log fold change) = {0.890201 (a), −0.932102 (b), 0.674794 (a)}

<u>Panel</u> ‘Only normal in zone’ (3 genes)

Same as panel with ‘only normal above’.

##### Cancer type

KIRP (32 normal samples, 289 tumor samples)

<u>Panel</u> ‘Only tumor above’ (3 genes)

*HK2*, *AC124798.1* and *NME1*

Reference (FPKM) = {0.89216, 1.33416, 4.97076}

Threshold (log fold change) = {1.55936, 0.760797, 0.652767}

<u>Panel</u> ‘Only normal above’ (1 gene)

*UMOD*

Reference (FPKM) = 1882.42

Threshold (log fold change) = −5.42522

<u>Panel</u> ‘Only tumor below’ (2 genes)

*WLS* and *MFSD4A*

Reference (FPKM) = {39.3454, 47.4444}

Threshold (log fold change) = {−0.561201, −2.64825}

<u>Panel</u> ‘Only tumor outside’ (3 genes)

*RGS12, ALG1* and *IFI27*

Reference (FPKM) = {3.17738, 4.89799, 10.6382}

Thresholds (log fold change) = {{−0.501801, 0.461124}, {−0.420365, 0.204626}, {−0.805476, 1.21677}}

<u>Panel</u> ‘Only tumor inside’ (6 genes)

*RPS4Y1, AC092868.1, LINC00239, LILRB2, PTPRH, MT-TP* and *KLRK1-AS1*

Reference (FPKM) = {10.1287, 0.939402, 0.365015, 0.654664, 0.193983, 957.896, 0.423354} Thresholds (log fold change) = {{−3.92341, 2.24955}, {1.15676, 2.93682}, {1.24479, 3.0379}, {0.66605, 1.96766}, {1.63273, 4.20457}, {0.486656, 1.7337}, {1.49459, 2.5866}}

<u>Panel</u> ‘Only tumor in zone’ (2 genes)

Same as panel with ‘only tumor below’.

<u>Panel</u> ‘Only normal in zone’ (1 gene)

Same as panel with ‘only normal above’.

##### Cancer type

LUAD (59 normal samples, 535 tumor samples)

<u>Panel</u> ‘Only tumor above’ (3 genes)

*ALDH18A1*, *TRIM27* and *PYCR1*

Reference (FPKM) = {9.25462, 6.24107, 2.89561}

Threshold (log fold change) = {0.745953, 0.311544, 1.54081}

<u>Panel</u> ‘Only normal above’ (5 genes)

*STX11*, *CALCRL*, *SFTPC*, *FHL1* and *EDNRB*

Reference (FPKM) = {26.2114, 29.2525, 7522.07, 54.2349, 30.3858}

Threshold (log fold change) = {−0.243529, 0.0562003, 0.0695837, 0.056593, 0.10761}

<u>Panel</u> ‘Only tumor below’ (3 genes)

*EMP2, TMEM220* and *PPP1R11*

Reference (FPKM) = {170.492, 3.42656, 29.2238}

Threshold (log fold change) = {−1.25461, −0.46466, −0.219425}

<u>Panel</u> ‘Only tumor outside’ (4 genes)

*BEX4, QPRT, PLAC8* and *DNAJC4*

Reference (FPKM) = {37.122, 4.5182, 9.53153, 9.56566}

Thresholds (log fold change) = {{−0.396451, 0.321376}, {−0.971489, 0.794282},

{−1.30566, 1.13451}, {−0.75676, 0.536434}}

<u>Panel</u> ‘Only tumor inside’ (8 genes)

*NLN*, *MMP11, MYBL2, ENSG00000236850, SQSTM1, IGLV3-25, CHAD* and *SGPP2*

Reference (FPKM) = {1.34552, 0.357485, 0.950542, 1.20662, 34.4058, 28.9682, 0.70378, 7.46544}

Thresholds (log fold change) = {{0.747499, 2.08854}, {2.80669, 5.05784},

{1.61175, 4.63111}, {0.766337, 2.69256}, {0.357981, 1.34473}, {2.84521, 6.62962}, {2.05955, 3.52922}, {1.62414, 3.60168}}

<u>Panel</u> ‘Only tumor in zone’ (3 genes)

*EMP2, TRIM27* and *ALDH18A1*

Reference (FPKM) = {170.492, 6.24107, 9.25462}

Threshold (log fold change) = {−1.25461 (b), 0.311544 (a), 0.745953 (a)}

<u>Panel</u> ‘Only normal in zone’ (5 genes)

Same as panel with ‘only normal above’.

##### Cancer type

LUSC (49 normal samples, 502 tumor samples)

<u>Panel</u> ‘Only tumor above’ (2 genes)

*MRGBP* and *F12*

Reference (FPKM) = {4.11491, 0.352769}

Threshold (log fold change) = {0.405015, 0.876742}

<u>Panel</u> ‘Only normal above’ (3 genes)

*EMP2*, *ESAM* and *GKN2*

Reference (FPKM) = {138.996, 58.6668, 26.7761}

Threshold (log fold change) = {−0.767637, −0.306594, −0.48887}

<u>Panel</u> ‘Only tumor below’ (2 genes)

*RAMP2* and *ADGRF5*

Reference (FPKM) = {68.6615, 71.9583}

Threshold (log fold change) = {−1.07069, −0.630642}

<u>Panel</u> ‘Only normal below’ (3 genes)

*SLC2A1*, *RFC4* and *PPP1R14BP3*

Reference (FPKM) = {3.59379, 1.90837, 8.07671}

Threshold (log fold change) = {0.646135, 0.305688, −0.0675405}

<u>Panel</u> ‘Only tumor outside’ (4 genes)

*CES1*, *BLMH*, *PTK6* and *EPHX1*

Reference (FPKM) = {76.8643, 6.21458, 2.85958, 116.526}

Thresholds (log fold change) = {{−1.05284, 1.01772}, {−0.534815, 0.575194}, {−1.13735, 0.730033}, {−0.687555, 1.04353}}

<u>Panel</u> ‘Only tumor inside’ (6 genes)

*AGMAT, SLC16A1, PTMAP4, CCDC18, TBX18* and *Z84485.1*

Reference (FPKM) = {0.309999, 2.69623, 1.45923, 0.558796, 0.155988, 0.243373}

Thresholds (log fold change) = {{1.11876, 3.01349}, {1.73048, 3.49603},

{0.677218, 2.53514}, {0.858494, 1.91101}, {2.52656, 4.70144}, {1.29957, 2.42192}}

<u>Panel</u> ‘Only tumor in zone’ (2 genes)

*MRGBP* and *ADGRF5*

Reference (FPKM) = {4.11491, 71.9583}

Threshold (log fold change) = {0.405015 (a), −0.630642 (b)}

<u>Panel</u> ‘Only normal in zone’

*EMP2* and *HDAC2*

Reference (FPKM) = {138.996, 3.98041}

Threshold (log fold change) = {−0.767637 (a), −0.00464922 (b)}

##### Cancer type

PRAD (52 normal samples, 499 tumor samples)

<u>Panel</u> ‘Only tumor above’ (9 genes)

*EPHA10, UCN, TMEM86A, ENSG00000258297, FRMPD3, BCAT2, TUT7, HOXA10-AS* and *SRMS*

Reference (FPKM) = {0.391225, 0.571144, 1.57228, 4.09354, 0.343019, 11.7103, 10.3469, 0.975031, 1.13588}

Threshold (log fold change) = {1.3373, 1.33742, 0.761516, 0.607416, 1.27324, 0.780582, 0.937288, 0.831813, 2.0766}

<u>Panel</u> ‘Only normal above’ (14 genes)

*SEPTIN10, CA14, DNAJB4, TES, LRRC37A7P, APOBEC3C, PARM1, AL512363.1, RNA5SP342, RCBTB2, ASS1P1, SLC18A2, TRMT1L* and *AC093536.1*

Reference (FPKM) = {16.8358, 2.11777, 7.18043, 29.5951, 0.965984, 30.6936, 62.5733, 0.446229, 0.150767, 8.53492, 0.327261, 1.31571, 4.89661, 0.500265}

Threshold (log fold change) = {0.426986, 0.8031, 0.812733, 0.669396, 0.913199, 0.883948, 0.62929, 0.925758, 2.24141, 0.328946, 1.5279, 2.78262, 0.488484, 1.77176}

<u>Panel</u> ‘Only tumor outside’ (12 genes)

*NCALD, SPATS2L, REX1BD, POLR1B, POLL, TLE1, FAM189A2, N4BP2L2, SUV39H1, MOSPD2, U2AF2* and *UBXN6*

Reference (FPKM) = {2.45813, 10.9487, 5.39723, 3.0657, 5.51475, 10.2287, 8.14368, 4.67076, 2.24326, 2.52638, 32.354, 30.2444}

Thresholds (log fold change) = {{−1.21066, 1.22599}, {−0.485148, 0.550506},

{−1.0208, 0.897549}, {−0.819029, 0.638888}, {−0.450603, 0.480065}, {−0.710702, 0.80299}, {−1.21858, 1.13504}, {−0.613803, 0.623638}, {−0.378771, 0.364548}, {−0.858434, 0.648279}, {−0.472689, 0.362606}, {−0.627057, 0.420475}}

<u>Panel</u> ‘Only tumor inside’ (11 genes)

*SNHG3, SAMD5, FHIT, PLA2G7, TPRN, NETO2, SLBP-DT, AL954705.1, SYP, AC099518.1 and MOSPD3*

Reference (FPKM) = {2.54932, 1.29739, 1.1174, 7.77365, 3.75213, 0.393405, 0.46288, 0.278518, 0.632042, 0.45783, 9.76841}

Thresholds (log fold change) = {{1.06334, 2.52785}, {1.0579, 3.74196}, {0.534765, 1.79877}, {1.27537, 2.72526}, {0.765739, 2.09259}, {0.610066, 1.85239}, {1.21917, 2.59009},

{1.106, 2.17853}, {0.626793, 3.20142}, {0.956875, 2.20259}, {0.550237, 1.7017}}

<u>Panel</u> ‘Only tumor in zone’ (13 genes)

*EPHA10, UCN, TMEM86A, NKAIN1, ENSG00000258297, REX1BD, AC099518.1* and *OAT*

Reference (FPKM) = {0.391225, 0.571144, 1.57228, 1.24333, 4.09354, 5.39723, 0.45783, 35.1421}

Threshold (log fold change) = {1.3373 (a), 1.33742 (a), 0.761516 (a), {1.56016, 4.23033} (i), 0.607416 (a), {−1.0208, 0.897549} (o), {0.956875, 2.20259} (i), −0.802215 (b)}

<u>Panel</u> ‘Only normal in zone’ (11 genes)

*SEPTIN10, CA14, DNAJB4, TES, LRRC37A7P, APOBEC3C, PARM1, AL512363.1, RNA5SP342, SNORD104, RCBTB2, TBC1D7* and *APOC1*

Reference (FPKM) = {16.8358, 2.11777, 7.18043, 29.5951, 0.965984, 30.6936, 62.5733, 0.446229, 0.150767, 4.33524, 8.53492, 1.466, 2.80483}

Threshold (log fold change) = {0.426986 (a), 0.8031 (a), 0.812733 (a), 0.669396 (a), 0.913199 (a), 0.883948 (a), 0.62929 (a), 0.925758 (a), 2.24141 (a), −2.30014 (b), 0.328946 (a), −0.592224 (b), −1.65552 (b)}

##### Cancer type

STAD (32 normal samples, 375 tumor samples)

<u>Panel</u> ‘Only tumor above’ (3 genes)

*MTHFD1L*, *ZNF146* and *SLC35E4*

Reference (FPKM) = {1.38861, 10.5207, 0.82564}

Threshold (log fold change) = {1.05193, 0.483634, 0.558899}

<u>Panel</u> ‘Only normal above’ (5 genes)

*IGHV3OR16-13, DPT, MALL, MT-TY* and *MYZAP*

Reference (FPKM) = {4.59954, 35.5918, 4.16596, 52.7855, 3.04359}

Threshold (log fold change) = {3.868, 1.27223, 3.21166, 2.67099, 0.691261}

<u>Panel</u> ‘Only tumor below’ (6 genes)

*CFD*, *ALDOC*, *METTL7A*, *RPS14*, *ME1* and *ATG4A*

Reference (FPKM) = {120.314, 11.5507, 35.0596, 149.246, 13.5885, 5.42812}

Threshold (log fold change) = {−2.4316, −1.15265, −1.11414, −0.495072, −1.39205, −0.559754}

<u>Panel</u> ‘Only normal below’ (9 genes)

*QSOX2, NUDCD1, ZBTB33, ENSG00000269899, BID, NOL10, MTERF3, DYNC1H1* and *AGO2*

Reference (FPKM) = {2.35163, 1.96061, 3.7481, 1.55122, 3.12532, 5.01874, 4.3212, 19.6871, 3.138}

Threshold (log fold change) = {−0.237129, −0.10108, −0.369973, −0.501522, −0.547937, −0.0695365, −0.312886, −1.02442, −0.879971}

<u>Panel</u> ‘Only tumor outside’ (7 genes)

*MACROH2A2, MEX3D, H2AJ, RAB13, AC005520.2, AC005912.1* and *PODXL2*

Reference (FPKM) = {7.21663, 8.52646, 15.7759, 21.3465, 1.86453, 60.1152, 6.64062}

Thresholds (log fold change) = {{−0.737553, 0.547616}, {−0.794033, 0.705834},

{−0.63225, 0.85081}, {−0.67364, 0.623539}, {−1.11841, 1.05993}, {−2.01781, 1.59006}, {−2.03466, 1.26447}}

<u>Panel</u> ‘Only tumor inside’ (5 genes)

*ZSWIM4, ESPL1, EFNA3, S1PR5* and *AL158206.1*

Reference (FPKM) = {2.69178, 1.16598, 1.0856, 0.381976, 8.57164}

Thresholds (log fold change) = {{0.44067, 2.0316}, {0.597465, 2.66237},

{0.339062, 3.14304}, {0.649493, 2.60641}, {−1.56043, 0.387396}}

<u>Panel</u> ‘Only tumor in zone’ (3 genes)

Same as panel with ‘only tumor above’.

<u>Panel</u> ‘Only normal in zone’ (5 genes)

*QSOX2, IGHV3OR16-13, DPT, XPO5* and *BID*

Reference (FPKM) = {2.35163, 4.59954, 35.5918, 3.18288, 3.12532}

Threshold (log fold change) = {−0.237129 (b), 3.868 (a), 1.27223 (a), −0.0716832 (b), −0.547937 (b)}

##### Cancer type

THCA (58 normal samples, 510 tumor samples)

<u>Panel</u> ‘Only tumor above’ (5 genes)

*METTL7B, GJC1, TYW1, SUSD4* and *ZMAT3*

Reference (FPKM) = {2.09007, 0.642125, 5.65877, 0.74864, 3.07059}

Threshold (log fold change) = {1.70463, 0.868901, 0.363749, 0.762627, 0.784466}

<u>Panel</u> ‘Only tumor below’ (8 genes)

*TFF3, TRMO, IGFBP6, AC106795.2, PRKACB, MPP1, RBPMS-AS1* and *DIO1*

Reference (FPKM) = {656.882, 5.66109, 22.096, 3.2275, 8.63334, 5.6294, 5.90389, 95.8447}

Threshold (log fold change) = {−2.10598, −0.396442, −1.72369, −1.14521, −0.618057, −0.383112, −0.910326, −2.74926}

<u>Panel</u> ‘Only tumor outside’ (4 genes)

*MXRA8, ORMDL3, ALDH4A1* and *FLT1*

Reference (FPKM) = {15.5592, 35.5455, 4.50101, 7.96758}

Threshold (log fold change) = {{−1.58551, 0.880012}, {−0.449449, 0.426211}, {−0.720023, 0.766703}, {−1.54347, 1.21369}}

<u>Panel</u> ‘Only tumor inside’ (6 genes)

*SERINC2, AC073343.1, ABCC6P2, GABRD, MMP11* and *FOLH1*

Reference (FPKM) = {12.5867, 3.16589, 0.365173, 0.434782, 0.332661, 0.238184}

Threshold (log fold change) = {{0.92633, 2.62982}, {−2.28176, −0.67791},

{1.03958, 2.4928}, {2.45567, 4.05592}, {1.55137, 2.55219}, {1.32925, 3.05291}}

<u>Panel</u> ‘Only tumor in zone’ (4 genes)

*METTL7B, CLU, NID1* and *TGFA*

Reference (FPKM) = {2.09007, 382.801, 4.08848, 3.3834}

Threshold (log fold change) = {1.70463 (a), {−0.87064, 0.778572} (o), 0.835444 (a), 0.877558 (a)}

<u>Panel</u> ‘Only normal in zone’ (13 genes)

*AC109326.1, AL034374.1, RIF1, BCL2L11, MMRN1, GPM6A, FNBP1, Z93241.1, UGT2B11, ENY2, ADH1B, OTUD7B* and *TRAJ58*

Reference (FPKM) = {2.20965, 3.68123, 3.7348, 11.0026, 7.66471, 3.80712, 15.4136, 0.894349, 0.566833, 8.20983, 3.71189, 3.58097, 0.125009}

Threshold (log fold change) = {0.385581 (a), 0.273349 (a), 0.619417 (a), 0.321295 (a), 0.898811 (a), 0.651311 (a), 0.62985 (a), 0.619176 (a), 0.733642 (a), 0.241412 (a), 1.2019 (a), {−1.465, 1.34669} (o), 2.4907 (a)}

##### Cancer type

UCEC (23 normal samples, 552 tumor samples)

<u>Panel</u> ‘Only tumor above’ (1 gene)

*TBC1D7*

Reference (FPKM) = 0.832865

Threshold (log fold change) = 0.577791

<u>Panel</u> ‘Only normal above’ (1 gene)

*PLSCR4*

Reference (FPKM) = 16.3218

Threshold (log fold change) = −0.971837

<u>Panel</u> ‘Only tumor below’ (2 gene)

*EZH1* and *TMEM245*

Reference (FPKM) = {13.5667, 18.0111}

Threshold (log fold change) = {−0.72488, −0.462326}

<u>Panel</u> ‘Only normal below’ (2 genes)

*TBC1D7* and *JPT1*

Reference (FPKM) = {0.832865, 5.19926}

Threshold (log fold change) = {0.577791, 0.767416}

<u>Panel</u> ‘Only tumor outside’ (3 genes)

*RNF10*, *TMEM101* and *PGK1*

Reference (FPKM) = {28.5136, 16.0865, 38.7264}

Thresholds (log fold change) = {{−0.113732, 0.1091}, {−1.10784, 0.986624},

{−0.5956, 0.257718}}

<u>Panel</u> ‘Only tumor inside’ (4 genes)

*HOXD10*, *MARCKSL1*, *RIPK2* and *SCD*

Reference (FPKM) = {2.98662, 93.5577, 4.6923, 4.59202}

Thresholds (log fold change) = {{−1.56749, 2.20744}, {0.70168, 2.50881}, {0.753443, 2.16763}, {1.8648, 3.47497}}

<u>Panel</u> ‘Only tumor in zone’ (1 gene)

Same as panel with ‘only tumor above’.

<u>Panel</u> ‘Only normal in zone’ (1 gene)

Same as panel with ‘only normal above’.

### B. Methodology

#### Data

TCGA holds a publicly accessible database of gene expression profiles coming from cohort studies with hundreds of normal tissue and solid tumor biopsy samples, classified by histopathological techniques (TCGA Research Network, 2006). The size of the data universe varies with the cancer type, and it is always skewed toward tumorous samples. Expression profiles are collected through RNA-Seq and register 60483 genes per sample (ibid.). TCGA reports gene expression values in the customary units of Fragments Per Kilobase of transcript per Million mapped reads (FPKM).

**Figure B1:**
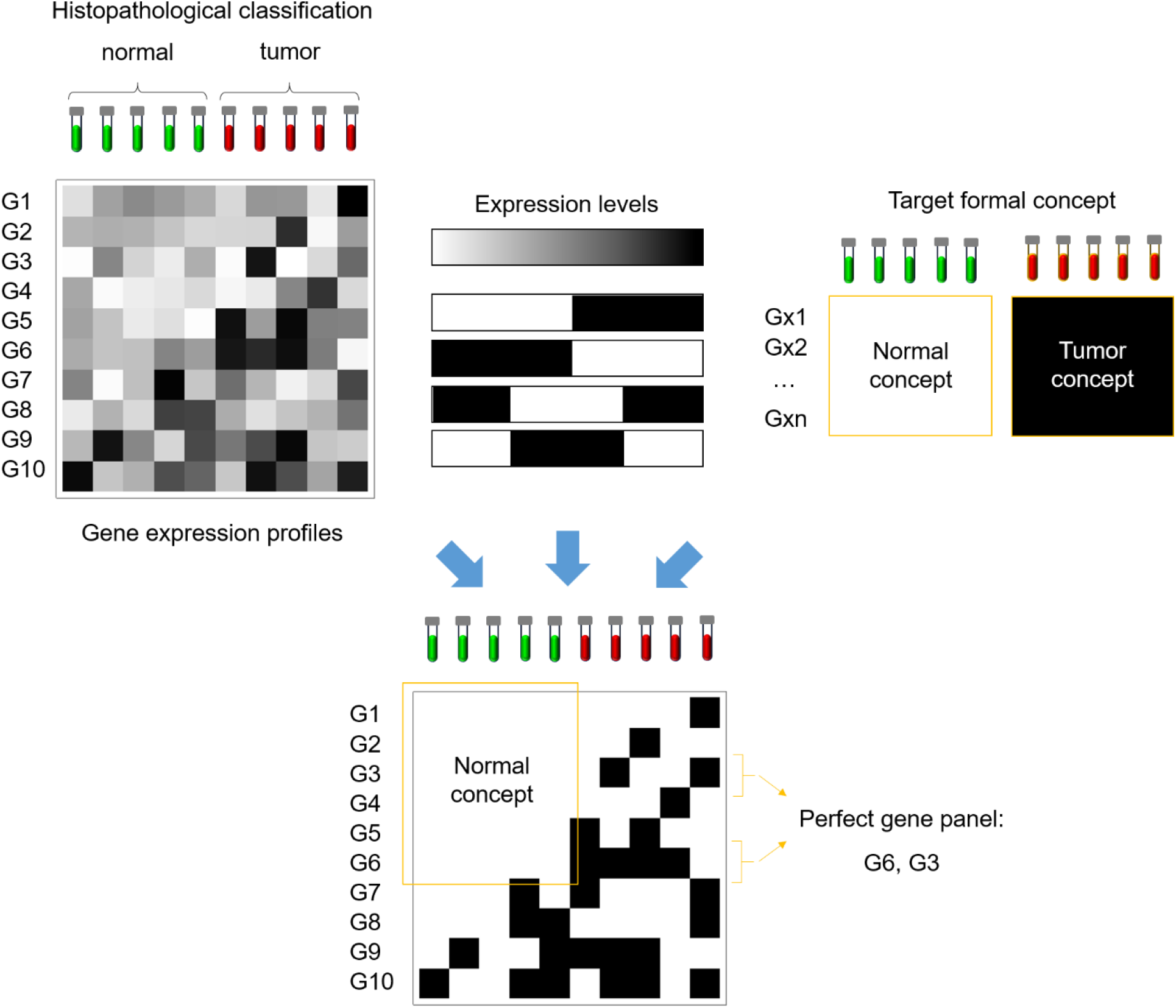
Schematics of our FCA / RST mining approach. In the upper panels, the main ingredients of the procedure are displayed. From left to right, those are (a) the gene expression profiles along with the histopathological classification of clinical samples, (b) all the dichotomizations of the gene expression levels we explored, and (c) the formal concepts from which the perfect panel can be built, based either on normal or tumor samples. The gene expression profiles (data to be mined) are extracted from the TCGA database, and afterwards minimally preprocessed by taking the fold change with respect to a normal reference. The schematics of (b) suggests that we associate normal-like or tumor-like expression levels (displayed in white and black) to different values of RNA-Seq counts that could be above threshold value, below it, outside a reference interval, or within it. In (c), we try to convey the essential nature of a formal concept, which is that they should be described by genes that take normal-like or tumor-like expression values for all normal samples or all tumor samples, respectively. In the lower section (d), we show the result of a mix of the three upper ingredients giving rise to a perfect gene panel. The example makes use of one of the possible dichotomizations (the first) and the normal concept. Notice that G6 and G3 are selected among the G1-G6 pool since they constitute a small panel that is able to classify perfectly all samples.

#### Preprocessing

The gene expression distribution is heavy-tailed, with many low-frequency outliers (e.g., Bengtsson, Ståhlberg, Rorsman and Kubista, 2005). Therefore, the average measure of choice for the gene expression is the geometric and not the arithmetic mean (ibid.). However, RNA-Seq is not accurate at detecting low expression values and may produce spurious null readings for genes that are almost silenced (Sha, Phan and Wang, 2015). Hence, in order to take the geometric mean as a characteristic level of expression over a population, we apply a harmless shift of 0.1 FPKM to all data.

Let *e*_*gs*_(*t*) be the resulting expression level of the gene *g* in the sample *s* taken from a tissue *t*. Our analysis is the same regardless of the tissue we apply it to. Moreover, does not involve mixing data from different tissues. Hence, we can safely drop the subscript (*t*) in the following. For each tissue, we estimate the homeostatic (or reference) gene expression level as the geometric mean of *e_gs_* over the set of normal samples *N*, i.e.

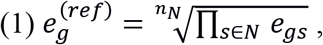

where *n_N_* is the number of normal samples.

Next, we take the log fold-change with respect to the latter reference,

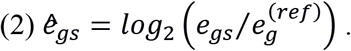

As a result, we treat up and down regulation symmetrically. *ê_gs_* is represented schematically as the gray-scale matrix in Figure B1 (a).

#### Dysregulation patterns

Now, we search for genes *g* matching specific dysregulation patterns. For differential expression cases, we have the following patterns:

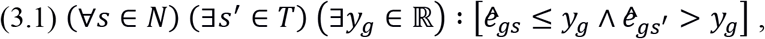

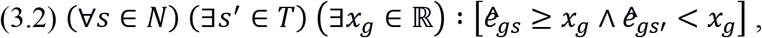

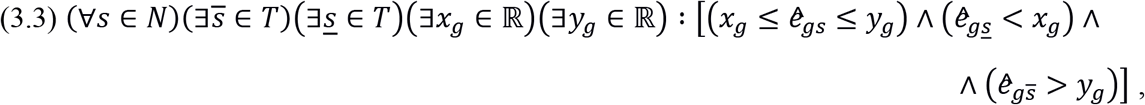

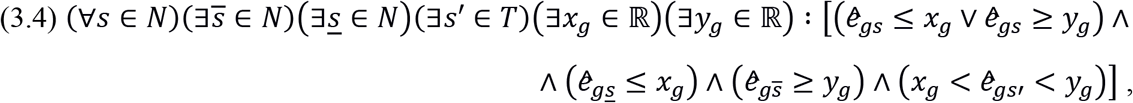

where *T* is the set of tumor samples. (3) correspond to genes with ‘only *tumor* above’, ‘below’, ‘outside’ and ‘inside’ in that order. Analogously, for the non-differential dysregulations:

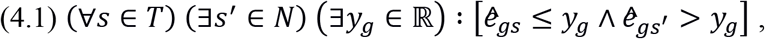

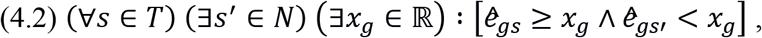

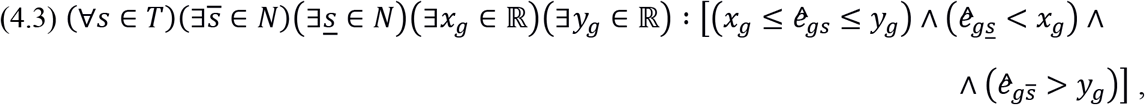

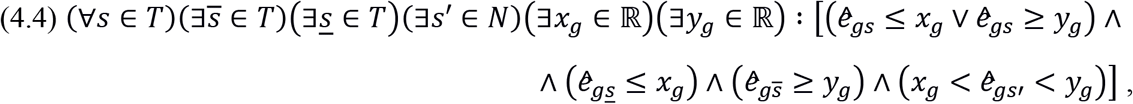

In passing from (3) to (4), we simply swapped the sets of normal and tumor samples. Conditions (4.1)-(4.4) correspond to genes with ‘only *normal* above’, ‘below’, ‘outside’ and ‘inside’ in that order. We represent (3) and (4) albeit schematically as the black and white patterned slabs in Figure B1 (b).

#### Expression thresholds

By setting the expression thresholds *x_g_* and *y_g_* to

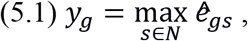

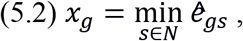

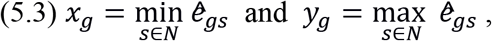

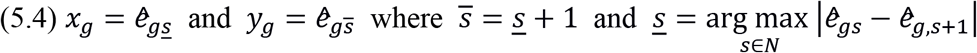

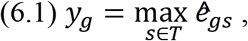

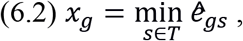

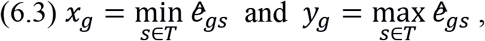

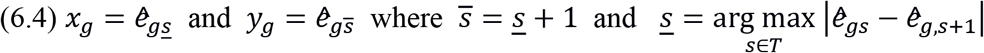

we automatically meet the quantification formulas over *s* in each case of (3)-(4), respectively. In (5.4) and (6.4), we suppose that normal and tumor samples (independently) are ordered in the order of their expression levels, *ê_gs_*. The rationale behind (5.4) and (6.4) is that the expression window without normal and tumor samples should be wide enough, for instance, with respect to any measure of the statistical dispersion of expression levels within the up and down regulated groups of normal and tumor samples, respectively. After (5)-(6), we only need to check

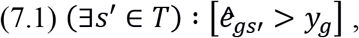

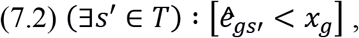

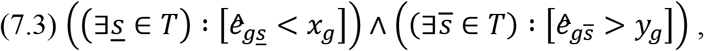

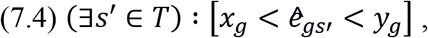

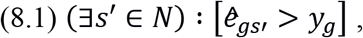

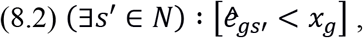

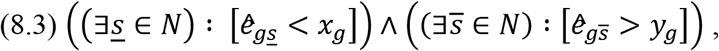

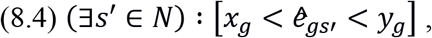

to assert that the gene *g* follows a particular dysregulation pattern. Conditions (7) correspond to (3)-(5) as (8) do to (4)-(6).

Thresholds reported in Section A of the Supplementary Information are computed as:

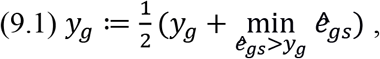

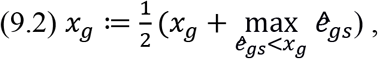

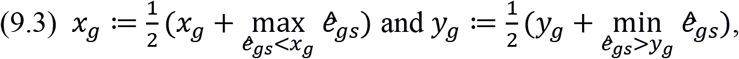

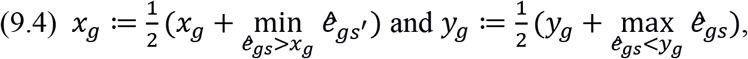

#### Statistically significant dysregulations

The check proposed in (7)-(8) may still come out true as a result of the statistical sampling, i.e., by chance. To select statistically significant genes, we demand that they pass a Fisher’s exact test for the classification of tumors and normal samples. When *g* exhibits a differential expression pattern (3.*n*), *A* is the set of samples for which (7.*n*) holds, assuming (5.*n*) [*n* within {1, 2, 3, 4}]. On the other hand, when *g* exhibits a non-differential dysregulation pattern (4.*n*), *A* is the set of samples for which (8.*n*) holds, assuming (6.*n*) [*n* within {1, 2, 3, 4}]. For differentially expressed genes, a successful Fisher test requires

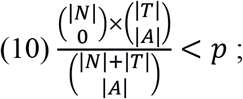

whereas for non-differentially dysregulated ones, we have

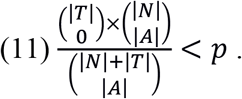

The cardinality of set *X* is denoted as |*X*|. Note that |*A*| corresponds to the number of *true* tumors for differentially expressed genes (3) and *true* normal samples for non-deferentially dysreg-ulated genes (4), respectively. In our account, dysregulated genes are the same as the classifier genes introduced in the Results (main text). That is, they do not produce false *tumors* in the case differentially expressed genes or false *normal* samples in the case of non-differentially dysregulated ones. Hence, the zero in the binomial coefficients of (10) and (11). We set a *p*-value of 1% in order to secure a high confidence level. Indeed, the latter choice means that only a 1% of the dysregulated genes we spot are incorrectly cataloged as dysregulated.

#### Special dysregulation patterns: only tumor/normal outside

Genes with ‘only tumor outside’ (3.3) and ‘only normal outside’ (4.3) are dysregulation cases where normal and tumor samples, respectively, fall within bounds whereas some samples from the complementary group lie both above and below them. The gene set meeting (3.3) partially overlaps with those sets for which either (3.1) or (3.2) hold. Similarly takes place for (4.3) and (4.1)-(4.2). The overlap is scaled down by demanding statistical significance in (10)-(11). However, this still leave us with ‘only tumor/normal outside’ genes that are already cataloged as ‘only tumor/normal above’ or ‘below’.

Besides the redundancy, the latter is indicative of a poor characterization of the dysregulation pattern in question, either (3.3) or (4.3). We want such patterns to be associated with commensurate amounts of samples below and above the expression thresholds in (5.3) or (6.3), i.e., for *ê_gs_* < *x_g_* and *ê_gs_* > *y_g_*. Imposing the previous constraint allows us to further reduce the overlap between the gene sets with ‘only tumor/normal outside’ and ‘only tumor/normal above’, as well as between those with ‘only tumor/normal outside’ and ‘only tumor/normal above’.

In practice, we set up one-tailed binomial tests for (3.3) and (4.3) dysregulations. The conditions of satisfaction for such test are

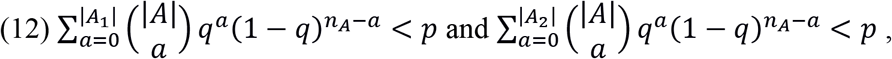

where *A_1_* and *A_2_* are subsets of *A* for which *ê_gs_* < *x_g_* and *ê_gs_* > *y_g_* obtain, respectively. Conditions (12) stipulate that the number of samples within *A_1_* and *A_2_* is such that the probability of finding an *A* sample either in *A_1_* or in *A_2_* cannot fall under *q*. A perfect balance between the number of samples in each group amounts to set *q* = 1/2. Our choice of *q* = 1/10 allows us to consider as well other less symmetric cases, such as a bimodal distribution with a distinctive anti-mode.

#### Special dysregulation patterns: only tumor/normal inside

Genes with ‘only tumor inside’ (3.4) and ‘only normal inside’ (4.4) are dysregulation cases where normal and tumor samples, respectively, lie outside an expression window whereas some samples from the complementary group fall within it. The gene set meeting (3.4) partially overlaps with those sets for which either (3.1)-(3.2) hold. Similarly takes place for (4.4) with (4.1)-(4.2). Again, the overlap is scaled down by requesting statistical significant dysregulations but it is not enough to consider them independent from each other.

We still require that such patterns to be associated with commensurate amounts of samples below and above the expression thresholds in (5.4) or (6.4), i.e., for *ê_gs_* < *x_g_* and *ê_gs_* > *y_g_*, which leads similar one-tailed binomial tests. For differentially expressed genes, we get

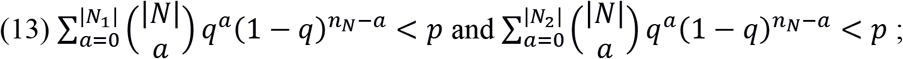

whereas for non-differentially dysregulated genes, we have instead

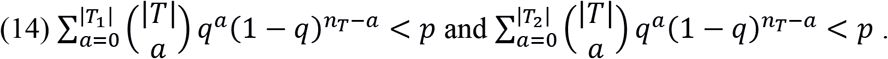

*N_1_* and *N_2_* are subsets of *N* for which *ê_gs_* < *x_g_* and *ê_gs_* > *y_g_* hold, respectively. Analogously, *T_1_* and *T_2_* are subsets of *T* for which *ê_gs_* < *x_g_* and *ê_gs_* > *y_g_* hold, respectively, and *n_T_* is the number of tumor samples. Conditions (13)-(14) admit a similar interpretation to those of (12).

Additionally, we demand that *y_g_* – *x_g_* > *δ*. In particular, we have set *δ* = 0.5 which prevents genes with unimodal expression distributions for normal samples (if differentially expressed) or tumor samples (if non-differentially dysregulated) to slide in. Note that, in the case of even log fold change distributions for differentially expressed genes, *y_g_* – *x_g_* > 0.5 contemplates customary normal bounds (namely, two and half-fold the reference expression level).

#### From gene expression profiles to a gene dysregulation checklist

We build up the binary matrix M = ||*m_gs_*|| with same number of samples *S* as the original data and all dysregulated genes *g* that are associated with a dysregulation pattern in (3)-(4). For a differential expression pattern (3.*n*), we set *m_gs_* = 1 if (15.*n*) obtains and otherwise *m_gs_* = 0, whereas for a non-differential dysregulation pattern (4.*n*), we set *m_gs_* = 0 if (14.*n*) obtains and otherwise *m_gs_* = 1. Condition (15.*n*) reads as follows

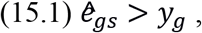

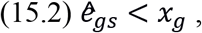

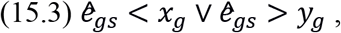

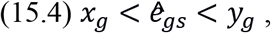

depending on the value of *n*. *x_g_* and *y_g_* values are defined in (5)-(6) for each case. A non-dysregulated gene *g* contains *m_gs_* = 0 for all samples *s*. Therefore, for a gene expression profile *s*, *m_gs_* is a checklist of all cancer-related dysregulations.

#### Formal concepts analysis: normal and tumor concepts

Let us establish minimal grounds for a formal concept analysis (FCA). First, we introduce the derivation operators (Lai and Zhang, 2009; Duntsch and Gediga, 2002) as

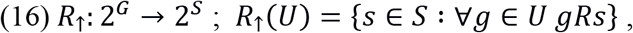

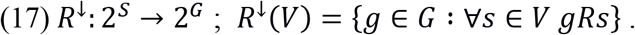

In the original formulation, *G* and *S* are sets of attributes and objects, respectively, whereas *gRs* is a logical condition stating that *g* is an attribute of the object *s*. The triple (*G,S,R*) is called a context.

In our context, *G* is the gene set (i.e., |*G*| = 60483), *S* the sample set (i.e., *S* = *N*∪*T*) and *gRs* a relation between the gene *g* and the sample *s*. Explicitly, if *g* conforms to a differential expression pattern, then *gRs* is equivalent to *m_gs_* = 0, meaning that the gene *g* is considered to be non-dysregulated in sample *s*. Else, if *g* conforms to a non-differential dysregulation pattern, then *gRs* is equivalent to *m_gs_* = 1, which means that the gene *g* is considered to be dysregulated in sample *s*.

A formal concept (definition) is a pair (*U,V*) with *V* = *R*_↟_(*U*) and *U* = *R*^↓^(*V*), where *U* is an attribute subset and *V*, an object subset. Therefore, for a formal concept we have *R*_↟_ (*R*^↓^(*V*)) = *V* or, equivalently, *R*_↟_ (*R*^↓^(*U*)) = *U*. In formal concepts: (a) every object in *V* has an attribute in *U*, (b) for every object outside *V*, there is at least one attribute in *U* that it does not have, and vice versa (c) for every attribute outside *U*, there is at least one object in *V* that does not have it.

A normal concept (definition) is a formal concept with *V* = *N*, whereas a tumor concept (definition) is a formal concept with *V* = *T*. Taking the set of differentially expressed genes 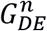 meeting (3.*n*), we have 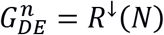. Similarly, for the set of non-differentially dysregulated genes 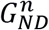 meeting (4.*n*), we get 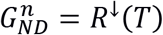. Indeed, from the definition of *m_gs_* in connection with (3) or (4), it follows that ∀*s* ∈ *N* (*m_gs_* = 0) or ∀*s* ∈ *T* (*m_gs_* = 1), respectively. However, neither 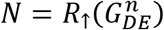 nor 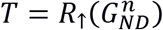 holds necessarily (i.e., by construction). Whether such conditions verify or not, it is to be checked in the data. Remarkably, for any subcategory of differential expression (3.*n*) and every cancer type in Table I, *N* = *R*_↟_(*G_DE_*) obtains (see Table II). The case is different for independent subcategories of non-differential dysregulations (4.*n*), where *T* = *R*_↟_(*G_ND_*) is not always verified for all cancer types under study. However, combining the four dysregulation patterns in (4), we still get tumor concepts for the cancer types in Table I.

In normal concepts, every tumor sample features at least a differential expression, and every normal sample is free from differential expressions. Therefore, normal concepts characterize tissue-specific homeostasis. On a causal interpretation, we expect that differentially expressing any given gene, may lead to cancer. Conversely, in tumor concepts, every normal sample exhibits a signature of normal regulation, and every tumor sample is free from normal regulations. Therefore, tumor concepts characterize cancer types. Here, the causal interpretation suggests that recovering the normal regulation pattern of a given non-differentially dysregulated gene, may kill a cancer cell or revert it to a normal status. Figure B1 (c) represents normal and tumor concepts by full-colored white or black genes-vs-samples rectangle with the sample side covering all normal or all tumors, respectively.

#### Rough set and reducts: perfect panels

Rough sets are usually defined by means of the following operators (Lai and Zhang, 2009; Duntsch and Gediga, 2002):

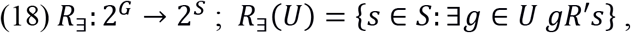

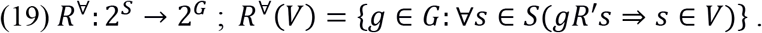

An *object-oriented* rough-set concept (definition) is a pair (*U,V*) with *V* = *R*_∃_(*U*) and *U* = *R*^∀^(*V*). In words, (i) all objects in *V* have at least an attribute in *U* and (ii) none of the objects outside *V* have an attribute in *U*.

Rough set theory (RST) is tightly related to FCA. In fact, (*U,V*) is an *object-oriented* rough-set concept in the context (*G,S,R*′) iff (*U,S*\*V*) is a formal concept in the complementary context *(G,S*,¬*R*), i.e., where the logical condition *gR*′*s* = ¬*gRs*. Henceforth, we take *gR*′*s* ≝ ′*gRs* with the interpretation of *gRs* given in the previous section.

The latter amounts to describe normal and tumor concepts also as rough-set concepts. Therefore, for a normal concept, we also have 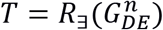 and 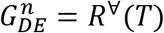, whereas for a tumor concept, we get 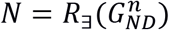and 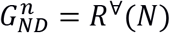.

A *formal-concept* reduct (definition) is (a) a subset of attributes from given formal concept with which the same formal concept is also reproduced, provided that (b) no attribute can be removed from the reduct without altering the formal concept. A formal concept may have many such reducts, and reducts corresponding to the same concept may vary in their membership and cardinality. Such reducts are not the *usual* reducts of RST (Pawlak, 1982). In fact, to the best of our knowledge, *formal-concept* reducts are here presented for the first time [cf. (Jia et al., 2016)]. However, more demanding notions in the same spirit have been already proposed by Zhang, Wei and Qi (2005).

Perfect panels (definition) are reducts of normal or tumor concepts, namely, gene subsets that preserve the normal or tumor concept and that cannot dispense of any gene member to do so. To build up panels, we carry out the following algorithm:

**Input:** Gene sets *G*′ and 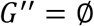
**Step 1:** Order *g* ∈ *G*′ by descending values of |*R*_∃_({*g*})| and update *G′*
**Step 2:** Take *G″’* = *G*’ \*G*” <u>keeping the order</u> in *G′*
**Step 3:** Collect in *G””* all *g* ∈ *G”’* producing a maximal value of |*R*_∃_(*G*′ ∪ *g*)| <u>keeping the order</u> in *G”’*
**Step 4:** Take the first *g* ∈ *G*”’ and update *G*” = *G*” ∪ *g*
**Step 5:** Iterate from **Step 2** until |*R*_∃_(*G*′)| = |*R*_∃_(*G*″)|
**Output:** Updated gene set *G”*

The latter produce *formal-concept* reducts if *G′* is the gene set of a normal or tumor concept. Note that the size of panels generated by the latter algorithm is not necessarily minimal, in the sense of being the smallest collection of genes achieving (a) and (b) [vide supra]. However, in practice, the size of the resulting panels ranges from 1-20 genes. Figure B1 (d) illustrates the process of finding a perfect panel from a formal concept.

When *G′* is not the gene set of a normal [tumor] concept but matches *G*′ = *R*^∀^(*N*) [*G*′ = *R*^∀^(*T*)] for at least a pattern of gene dysregulations (3) [(4)], the above algorithm produce a gene set that is able to cast *combinatorically* as much tumor [normal] samples as the full set of genes *G′* (see **Step 5**). Other kinds of gene subsets *G*′ are not suitable as input for the former algorithm.

#### Robustness of our findings

Both normal and tumor concepts are sensitive to variations in the universe of samples. In fact, a previously nonexistent (or statistically insignificant) exclusion window for a kind of samples – either normal or tumorous– may open up (or become statistically significant) as a result of an increase of the complementary kind or a decrease of that same kind of samples. And, vice versa: an existing (or statistically significant) exclusion window for a kind of samples may shut down (or become statistically insignificant) as a result of an increase of that same kind or a decrease of the complementary kind of samples. Therefore, varying the number of any kind of samples may affect the gene membership of both normal and tumor concepts

As anticipated, this issue is less pressing for a normal concept than it is for a tumor concept. The reason is twofold. One way a normal or a tumor concept could be shrunk is by discarding tumor or normal samples, respectively. But since the membership of normal concepts is more demanding on the number of true tumors than the membership of tumor concepts is on the number of true normals (see above), we expect that the normal concept to be less affected upon discarding tumor samples than the tumor concept should be upon discarding normal samples. Ultimately, this is a contingent fact depending on the great difference between the number of tumor and normal samples within the TCGA data for any cancer type.

The other way in which a normal or a tumor concept could be shrunk is through the incorporation of further normal or tumor samples, respectively. Here, the argument comes from a different corner. González et al. (2022) have noted a difference in variance displayed by both the homeostatic and cancer “attractor” states (Huang, Ernberg and Kauffman, 2009) in the gene expression landscape. In fact, the number of available microstates for cancer is several orders of magnitude above its counterpart for homeostasis (González, Quintela, León, Bringas Vega and Valdés Sosa, 2022). Hence, we do not expect to enhance our description of the homeostatic attractor upon an increase of normal samples as much as we expect to enhance our description of the cancer attractor upon an increase of tumor samples.

As normal and tumor concepts are sensitive to variations in the universe of samples, so too is any gene panel representing them. Indeed, gene panels are special subsets of concepts which cannot lose a gene member without compromising a perfect classification of samples. Therefore, depriving a concept from some of their genes may impact the perfect panel membership or worse, its sheer existence as a perfect classification scheme.

In order to assess the robustness of our results, especially in what regards the gene panels, we examine the *tumor* concept called ‘Only normal in zone’ for PRAD. Given that the tumor concepts are the more sensitive to a variation in the universe of samples, the ensuing analysis will suffice also to establish lower bounds for the robustness of our findings in normal concepts.

We quantify the robustness of our results with the following figures of merit: the sensitivity of the classification scheme based on the new data panel, the part of the original data panel that appears in the new data panel, and the part of the new data panel that appears in the original data panel. By choice, the new data panel is built up from those members of the original data panel that also entail a 100% specificity within the new data. It is worth noting that our null hypothesis for tumor concept genes is that they are non-differentially dysregulated (hence determining a tumor sample, whether true or false), whereas the alternative hypothesis is that they are normally regulated (hence determining a *true* normal sample). By ‘new data’ we mean the ‘original data’ minus normal or plus tumor samples, depending on the case. We generate further tumor samples by a process of bootstrapping. To wit, for each gene we perform the Kernel Density Estimation and resample the resulting probability density as many times as there are tumor samples in the original data. Normal and tumor samples (whether actual or simulated) are organized in decreasing order of the total number of gene dysregulations they exhibit within the original data. We take cumulative percentages of the more dysregulated samples in order to link variations in the universe of samples with any of our figures of merit.

We also refer to the former ‘new data’ as the ‘training set’. The term is borrowed from machine learning, where it represents that part of the data the algorithms learn from. In the analysis with respect to a decrease of actual normal samples, our training set is composed of a subset of normal samples and the full set of actual tumor samples, whereas in the analysis with respect to an increase in tumor samples, it comprises the full set of actual samples (normal and tumorous) plus a subset of simulated tumor samples. The notion of ‘training set’ is usually opposed to that of a ‘validation set’, which is the part of the data that is used to assess the quality of the learning from the training set. In the analysis with respect to a decrease of actual normal samples, our validation set consists of the remaining normal samples, whereas in the analysis with respect to an increase in tumor samples, it includes the remaining, simulated tumor samples. The former and latter sets allow us to validate the corresponding sensitivity and specificity of the new data panel.

Figure B2 conveys the results of our robustness analysis. We plot all figures of merit against an increasing number of normal (left column) and additional tumor samples (right column). The top row displays the sensitivity and/or specificity as function of the new data universe, whereas the bottom row shows how much gene panel membership is affected by it. Notice that we get 100% specificity by definition of the tumor concept within the training set. Additionally, note that when the validation set is without normal or tumor samples it is meaningless to assign to it any sensitivity or specificity, respectively.

We find that the sensitivity ramps up quickly for the training and validation sets of the left column, reaching 100% around 70% of the normal samples, and that it is always optimal for the training set of the right column, irrespective of the number of additional tumor samples (see Fig. B2, top row). On the other hand, the specificity fluctuates around 95% upon increasing tumor samples (see Fig. B2, top right). Moreover, we observe that both the contribution of the original data panel to the new data panel and vice versa approach its maximal near the limit of the whole set of actual samples. In particular, the panel seems to undergo a minor replacement of just one of the original genes by two new genes near 70% of the normal samples (left column). Remarkably, the original panel falls completely within the new data panel even when supplying up to an extra 50% of the tumor samples. In the latter range, the original panel gets completed by 6 new genes. Taken together all these features show the robustness of our mining scheme and the stability of the resulting gene panels.

**Figure B2:**
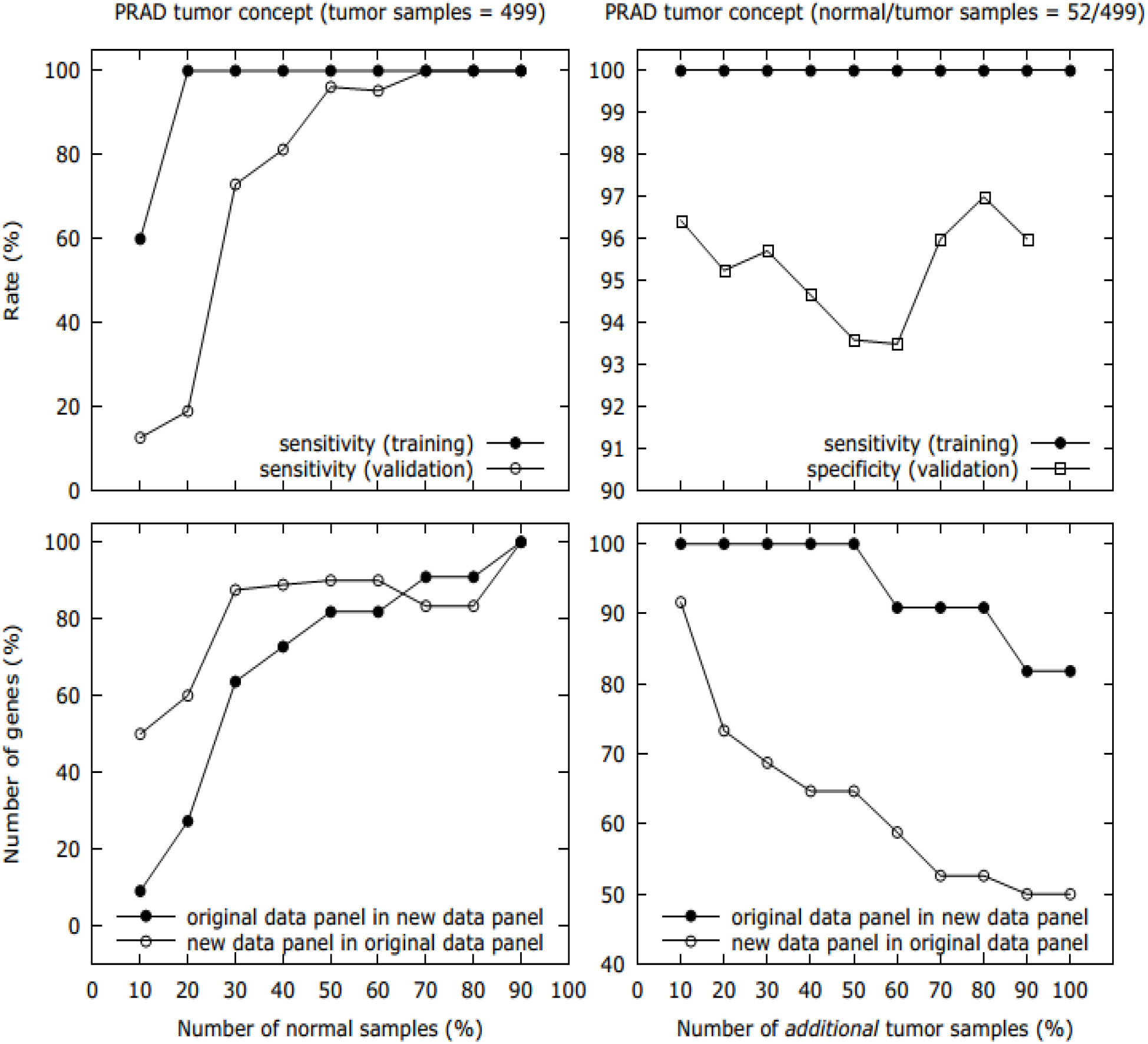
Robustness analysis for a PRAD tumor concept. In the left column, the number of actual normal samples taken into account is modified from 5 to 46 (out of 52), while considering all actual tumor samples (499) and only those. In the right column, actual data is progressively supplemented with further tumor samples (from 49 to 499) generated by a bootstrapping procedure. The maximum number of additional and actual tumor samples match. Samples are arranged in decreasing order of the number of dysregulations, and are selected cumulatively, from the most to the least dysregulated samples. At the top and where applicable, we plot the sensitivity (circles) and/or specificity (squares) rates for the new data panel within the training (filled symbol) and/or validation (empty symbol) sets. At the bottom, we show the part of the original data panel that appears in the new data panel (filled circles) and vice versa, the part of the new data panel that appears in the original data panel (empty circles).

#### Constraining the search

Hitherto, we have considered a principled way of searching for genes that match specific dysregulation patterns. We call it the ‘base method’. Next, we provide a stringent alternative for such a search according to which dysregulation in differentially expressed genes or normal regulation in non-differentially dysregulated genes conforms with accurate experimental detection by RNA-Seq standards. In particular, we require instances of dysregulation in a differentially expressed gene, as well as instances of normal regulation in a non-differentially dysregulated gene, to be associated with a gene expression value above 0.5 FPKM. Notice that reassessing cases of gene dysregulation in such a way may entail a shift in the expression thresholds defined before (see *Expression thresholds)* for panels ‘only tumor/normal above’ and ‘only tumor/normal inside’. In the case of panels with ‘only tumors/normals inside’, we further demand that the gene expression exclusion interval for normals/tumors be as wide as a unit of binary fold change (cf. *Special dysregulation patterns: only tumor/normal inside*).

Overall, we refer to the latter strategy as the ‘stringent method’, as opposed to the previously introduced ‘base method’. Results for the base and stringent methods are shown in Sections A1 and A2. First, note that, in some of the panels that stem from the base method, there are genes that are supposed to classify beyond low-expression detection limits (i.e., 0.5 FPKM). For example, the base method attributes *APLN* the potential for true classification of normal samples below 0.08 FPKM in the case of LIHC (see Section A1). This is systematically avoided in the stringent search by construction.

The stringent method does not always succeed in building up perfect gene panels there where the base method does. For instance, according to the stringent criterion, we cannot assemble a panel with ‘only tumor below’ that produces perfect classifications of prostate samples (cf. Section A1). Nevertheless, following either method, we can create comprehensive panels with ‘only tumor/normal in zone’ for all cancer types.

When successful at building a perfect panel, the stringent method may end up adding or subtracting genes from the analogous base-method panel. However, the eventual resizing is typically limited to an increase of one or two panel members. For instance, the panel with ‘only tumor above’ from PRAD is composed of either 8 or 9 genes depending on whether it was generated by the base or the stringent method. Remarkably, there are a couple of panels that get shrink when switching to the stringent method, as it is the case of the ‘only tumor below’ from LUAD, with 4 and 3 genes in the base and stringent method, respectively. Perhaps the fact that we robustly obtain small-sized panels upon moving to a more stringent method could be seen as a signature of the low-dimensional boundary between cancer and homeostasis attractors within the multivariate gene expression landscape (Nieves and González, 2021).

Figure B3 shows the results of our robustness analysis in the case of the stringent method (cf. Fig. B2). Apart from quantitative details and noise at low number of normal samples and at low number of *extra* tumor samples, Fig. B2 and B3 are alike in every important respect.

**Figure B3:**
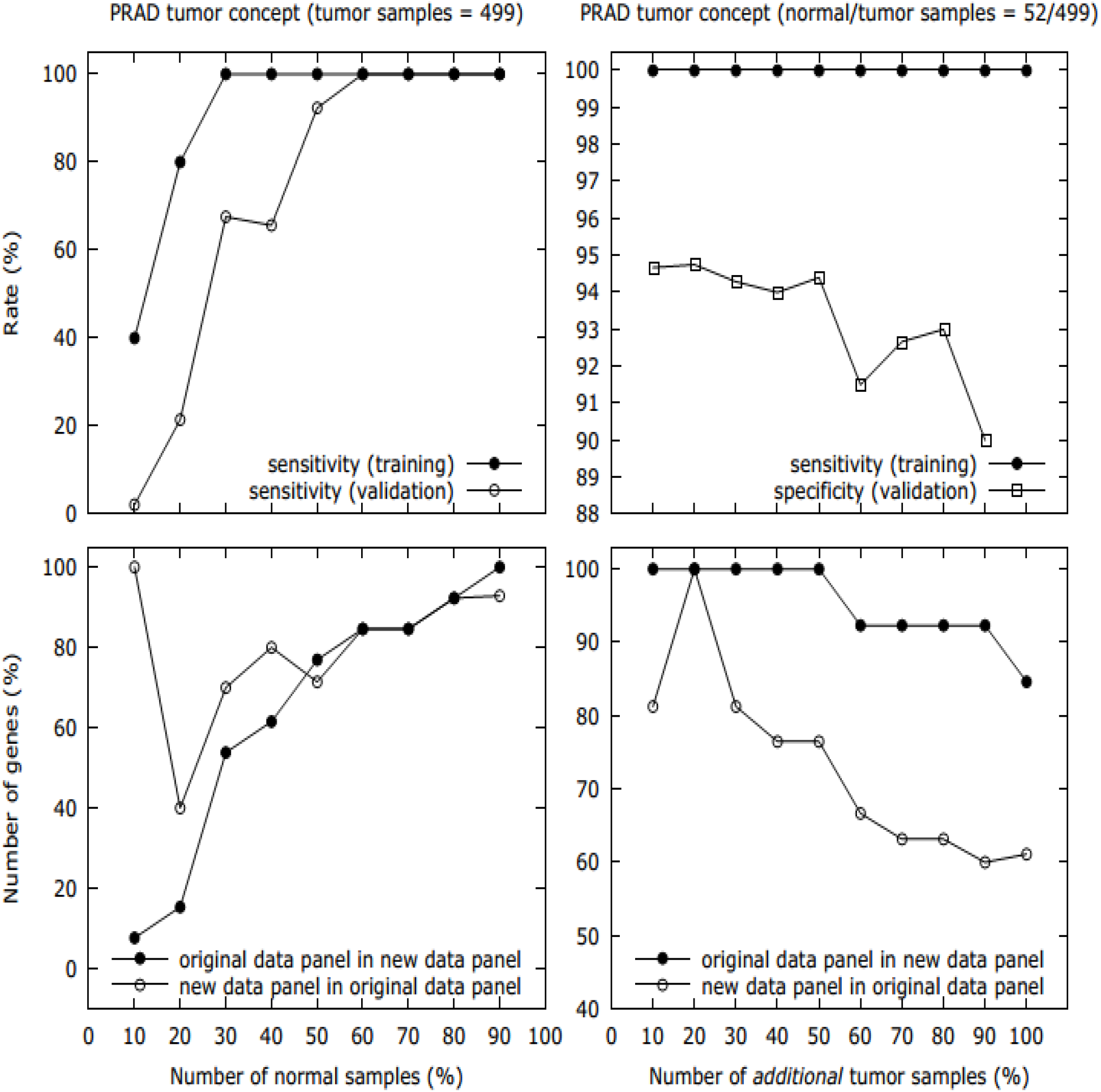
Same as Fig. B2 (base method) in the case of the stringent method. Figs. B2 and B3 are alike in all the key aspects tackled in the previous section.

#### Code and Repository URL

Our methodology is implemented in our home-made and self-standing Wolfram Mathematica code GenePan, which is hosted in the GitHub repository https://github.com/gabriel-gil/GenePan together with the relevant TCGA data for a prototypical run.

### C. Footnotes to the main text

C1: Our data is a collection of gene expression profiles, one per clinical sample. To avoid confusion between clinical and statistical sample, we will refer to ‘data universe’ when talking about the latter.

C2: We use kernel density estimation with a Gaussian kernel and the Silverman rule of thumb for the bandwidth [Silverman, B.W. (1986). *Density Estimation for Statistics and Data Analysis*. London: Chapman & Hall/CRC. p. 45. ISBN 978-0-412-24620-3].

C3: Assuming ergodicity of the gene expression, we can take profiles from different patients at the same time to be representatives of the evolution of the profile from a single patient over time. With such caveat, we can draw inferences about oscillations from the shape of single-gene expression distributions. Modes in the oscillatory coordinate are thus associated with turning points of oscillations. When modes are of different height, the oscillatory system linger around one of the turning points, matching the typical description of a pulse.

C4: As customary in Statistical hypothesis testing, type I errors are those incorrect rejections of the null hypothesis. Notice that these are also known as false positives. However, in our case the latter terminology might be misleading, since for us the null hypothesis can be either that the sample is normal or that it is tumorous, depending on the gene in question. We prefer not to call them false positives to avoid the interpretation that type I errors always refer to false tumors. Indeed, our type I errors can be false tumors or false normal samples for different genes.

C5: The non-overlapping expression window may be disconnected, as in the case where both up and down regulation lead to cancer, or in the case where only normal expression is oscillatory (see *Special dysregulation patterns: only tumor/normal outside* and …*inside* in Section B).

C6: There are two hypothesis tests interlinked in this analysis: one inherent to the classification of samples into normal and tumor, another related to the classification of genes into significant and non-significant. The former checks the expression of a certain gene in a sample in order to accept or reject the null hypothesis of it being a case, e.g., of a tumor (see *From gene expression profiles to a gene dysregulation checklist* in Section B). The p-value of such a classification is 0% for the chosen genes, meaning that the rejection of null hypothesis is infallible (no type I errors). The other hypothesis test (in this case, a Fisher’s exact test) proceeds by checking how many, e.g., true normal cases are classified by a gene in order to accept or reject the null hypothesis of it being a random (i.e., non-significant) classifier (see *Statistically significant dysregulations* in Section B). Here, we set an upper bound for the p-value to 1%, meaning that we allow up to 1% of the selected genes to be misclassified as significant. Other fine-grain significance criteria are adopted there where the exclusion zone is an interval or its complement (see *Specialdys regulation patterns: only tumor/normal outside* and …*inside* in Section B).

C7: Types II errors are cases of incorrect acceptance of the null hypothesis, also known as false negatives. We prefer not to use the latter terminology in order to avoid confusion. Here, type II errors can be either false normal or false tumor samples, depending on the gene. See also C4.

C8: We use a heuristic algorithm consisting in the iterative gene selection from a pool of gene classifiers and according to two criteria: (i) first and foremost, the next gene must be that which provides the best possible classification among the yet unselected genes, and (ii) secondarily, the next gene must be as redundant as possible with previously selected genes (see *Rough set and reducts: perfect panels* in Section B). Such a heuristic allows us to find small panels, in general less than 10 genes and in any case no greater than 19.

C9: The perfect panel size is minimal in the sense that removing any member leads to imperfect classifications (see *Rough set and reducts: perfect panels* in Section B). Note that there might be many different perfect panels for any cancer type, and indeed, some of them might be even smaller than our choice. However, looking for them goes beyond our main goal which is to find a set of combinatorial biomarkers for realistic diagnostics and therapy.

C10: Since we do not work with single-cell RNA-Seq data, the tumor-concept panel may include as well genes that are consistently activated in normal tissue-resident or immune cells in response to the presence of tumor cells within the environment.

